# An Alzheimer’s disease-associated common regulatory variant in a PTK2B intron alters microglial function

**DOI:** 10.1101/2023.11.04.565613

**Authors:** Erica Bello, Kathleen Long, Sho Iwama, Juliette Steer, Sarah Cooper, Kaur Alasoo, Natsuhiko Kumasaka, Jeremy Schwartzentruber, Nikolaos I Panousis, Andrew Bassett

## Abstract

Genome-wide association studies (GWAS) are revealing an ever-growing number of genetic associations with disease, but identifying and functionally validating the causal variants underlying these associations is very challenging and has only been done for a vanishingly small number of variants. Here we validate a single nucleotide polymorphism (SNP) associated with an increased risk of Alzheimer’s disease (AD) in an intronic enhancer of the *PTK2B* gene, by engineering it into human induced pluripotent stem cells (hiPSCs). Upon differentiation to macrophages and microglia, this variant shows effects on chromatin accessibility of the enhancer and increased binding of the transcription factor CEBPB but only subtle effects on PTK2B or CLU expression. Nevertheless, this variant results in global changes to the transcriptome and phenotype of these cells. Expression of interferon gamma responsive genes including chemokine transcripts and their protein products are altered, and chemotaxis of the resulting microglial cells is affected. This variant thus causes disease-relevant transcriptomic and phenotypic changes, and we propose that it acts by altering microglia reactivity, consistent with the role of these cells in progression of AD.

## Introduction

Alzheimer’s disease (AD) is the most common form of dementia, and has a strong genetic component estimated to be 60-80%[1]. AD is characterised by the presence of β-amyloid-containing plaques (Aβ) and intracellular neurofibrillary tangles of hyperphosphorylated tau, accompanied by neuroinflammation, neuronal, and synapse loss[2]. Multiple GWAS studies have collectively identified more than 80 genetic loci associated with disease[3–7], the vast majority of which do not affect protein coding sequence. Due to linkage disequilibrium at these loci, it is difficult to identify the causal variants. Multiple computational methods have been developed to prioritise variants that cause disease including statistical fine mapping, co-localisation with expression or chromatin accessibility quantitative trait loci (eQTL, caQTL), overlap with enhancer regions identified by for example the assay for transposase accessible chromatin (ATAC) or transcription factor binding in disease relevant cell types. However, none of these methods are able to unambiguously assign causal variants and, given the very small number of truly validated variants, it is difficult to benchmark their effectiveness. Experimental methods have also been developed to identify or prioritise variants such as massively parallel reporter assays (MPRA[8]) and genome engineering-based interrogation of enhancers (GenIE[9], GenIE-ATAC[10]) but these are limited to understanding the effects of variants on transcription or chromatin accessibility and do not directly demonstrate the downstream effects of these polymorphisms. Genome engineering of isogenic human cell models, such as differentiated hiPSCs differing in a single variant, provide an attractive means for understanding the role of specific SNPs on cell function and have been successful in understanding the role of strong effect familial mutations[11,12,13]. However, such systems have yet to be successfully applied to more than a handful of common genetic variants[14,15,16] and their effects are often confounded by variation between edited clones and during differentiation[17].

Neuroinflammation has been shown to be a crucial factor in many neurodegenerative disorders, especially AD[18]. Microglia are resident immune cells of the central nervous system and perform various functions under both homeostatic and disease conditions [19]. They play important roles in synaptic remodelling, synaptic pruning and phagocytosis of dead neurons[20]. Microglia have been implicated in AD by human genetics studies because of specific expression of AD-related genes in these cells[21–23], colocalisation of microglial eQTLs with AD GWAS hits[24,25] and implication of rare causal variants in microglial-specific genes, such as TREM2[26] and CD33[26,27]. Nevertheless, the exact role of microglia in AD pathogenesis is still unclear. Aβ and tau aggregates are able to activate microglia, stimulate cytokines and chemokine production and elicit an inflammatory response that can have two functions: it can be neuroprotective by increasing Aβ or tau clearance, but it can also contribute to Aβ and tau production and spreading and induce neurodegeneration and synaptic loss[28,29]. Microglia are known to be involved in clearance of aggregated proteins such as Aβ[30] and accumulation of extracellular Aβ is accelerated by insufficient microglial activation or phagocytic capacity[31]. However, depletion of microglia in amyloid precursor protein-transgenic mouse models was found to have little effect on plaque formation or maintenance. Microglia have also been found to potentially contribute to tauopathy by spreading tau protein via exosomes across brain regions[32] as well as contributing to its uptake and degradation[29]. Moreover, sustained microglia activation appears to contribute to neurodegeneration by impacting synaptic plasticity and causing neuronal damage[33,34]. As well as microglia, other myeloid cell types, such as infiltrating monocytes[35,36], choroid plexus[37] and perivascular macrophages[38–40], have also been found to play a role in AD pathogenesis. Circulating monocytes can infiltrate the brain and target cerebral Aβ deposits to reduce plaque load[41–43] and depletion of perivascular macrophages in the TgCRND8 mouse model of AD significantly increases vascular Aβ levels[44]. Conversely, β amyloid deposition in cortical blood vessels is reduced by stimulation of perivascular macrophage turnover[44]. Moreover, studies have shown that macrophages isolated from individuals with AD are poorly phagocytic for Aβ, more susceptible to apoptosis and have impaired chemotaxis[45,46].

Chemokines are produced by several cells in the central nervous system such as neurons, astrocytes, immune cells and particularly microglia. They are known to recruit peripheral immune cells through the blood brain barrier as well as activate brain resident macrophages and microglia[47]. Elevated levels of several chemokines have been found in plasma, cerebrospinal fluid and brain tissue of AD patients[48]. Proinflammatory chemokines are believed to contribute to neuroinflammation and chronic activation of microglia and macrophages in late and symptomatic stages of AD[47] and may constitute a useful biomarker to monitor disease progression[49]. However, chemokine-mediated microglia recruitment and phagocytosis aids clearance of Aβ oligomers and knockout of some chemokines, such as CCL2 (and CCR2 receptor) in mouse models of AD results in accumulation of intracellular soluble Aβ oligomers and decline of cognitive function[50–52]. Aβ can stimulate human monocytes and microglia to produce chemokines, such as CXCL8, CCL2 and CCL3, which also play a role in leukocyte migration to sites of inflammation[53].

PTK2B is a non-receptor tyrosine kinase that is expressed in neurons, microglia, astrocytes, monocytes and tissue-resident macrophages, and has diverse physiological and pathological roles including cell adhesion, cell migration, inflammatory responses, tumour invasiveness[54], neuronal development and plasticity[55,56]. PTK2B is a member of the focal adhesion kinase (FAK) subfamily and after autophosphorylation it recruits Src-family kinases[57] and interacts with multiple partners to activate various downstream signalling pathways, such as the MAPK/ERK pathway[55]. In neurons, PTK2B is implicated in synaptic plasticity[58] and neurite outgrowth[59]. In non-neuronal cells, such as osteoclasts[60], macrophages[61–63] and monocytes[64], PTK2B is involved in cell migration and regulation of the actin cytoskeleton downstream of integrins. In monocytes, the ROS-sensitive calcium channel TRPM2 can activate the PTK2B/ERK pathway, leading to nuclear translocation of nuclear factor-kB and increased chemokine production[65]. This pathway is activated by Aβ in microglia, where PTK2B is autophosphorylated and generates a positive feedback loop to further activate TRPM2[66]. PTK2B has been implicated in AD through a number of GWAS studies[4,6,67–74]. Evidence suggests that genetic deletion of Ptk2b rescues a number of Aβ-associated phenotypes, such as memory impairment, synapse loss and impaired synaptic plasticity in APP/PS1 animal models of AD[75]. Conversely, overexpression of Ptk2b has been reported to be protective in 5xFAD mice [76] and human neuronal systems [77]. Similarly, data collected from hemizygous PS19 (MAPT P301S, 1N4R) transgenic mice crossed with Ptk2b^−/−^ animals, points to a protective role for Ptk2b against Tau phosphorylation and Tau-induced behavioural deficits[78]. Thus, several lines of evidence point to PTK2B as an important player in the pathophysiology of AD but much remains to be learnt about the exact role of PTK2B in AD pathogenesis.

Clusterin (CLU) is a glycoprotein which is mainly secreted but also exists intracellularly, and is involved in a variety of cellular functions[79]. Secreted clusterin acts as a molecular chaperone which stabilizes misfolded proteins and can prevent protein aggregation[80]. CLU is also involved in modulating inflammatory proteins as well as proteins involved in cell survival and apoptosis[79]. Cellular stress can lead to an upregulation of CLU which activates anti-apoptotic pathways and enables cell survival[81–83]. CLU is expressed across many tissues, and in the brain it is expressed in neurons as well as astrocytes and microglia[79] . Secreted clusterin interacts and can bind Aβ, and evidence suggests that this can alter Aβ aggregation and promote clearance[81,84,85] These studies point to a neuroprotective role of CLU, however other evidence suggests that clusterin instead reduces Aβ clearance and increases neurotoxicity[86,87]. The CLU locus is the third greatest genetic risk factor for late onset AD and several SNPs in this gene alter AD susceptibility[79]. CLU is upregulated in the hippocampus, cortex and CSF of AD patients and it colocalises with Aβ [81,88].

AD GWAS studies have identified a locus on chromosome 8 which contains two independent association signals, one over PTK2B and one over CLU[67]. Mapping of chromosomal interactions at the PTK2B locus in microglia has shown that enhancers harbouring AD-risk variants interact with active promoters of both PTK2B and CLU[89] but further investigation is needed to identify causal variants at this locus and their putative target genes.

## Results

### Prioritisation of a putative Alzheimer’s disease variant at the PTK2B locus

GWAS have identified two independent genetic associations at the PTK2B-CLU locus[67]. The signal over the PTK2B gene colocalises with both an eQTL and caQTL in hiPSC-derived macrophages (Fig 1A)[90]. Re-analysis of publicly available datasets[25,90,91] shows that the risk haplotype is linked to both a downregulation of PTK2B gene expression in hiPSC-derived macrophages (Fig 1B and 1C) and an increase in chromatin accessibility at an enhancer within intron 6 of the PTK2B gene (Fig 1D). Interestingly, in this case there is a single variant in the credible set (rs28834970) that lies within the ATAC peak that is changed in accessibility in the caQTL analysis, strongly suggesting that this variant is causal (C being the risk allele).

**Figure 1.**
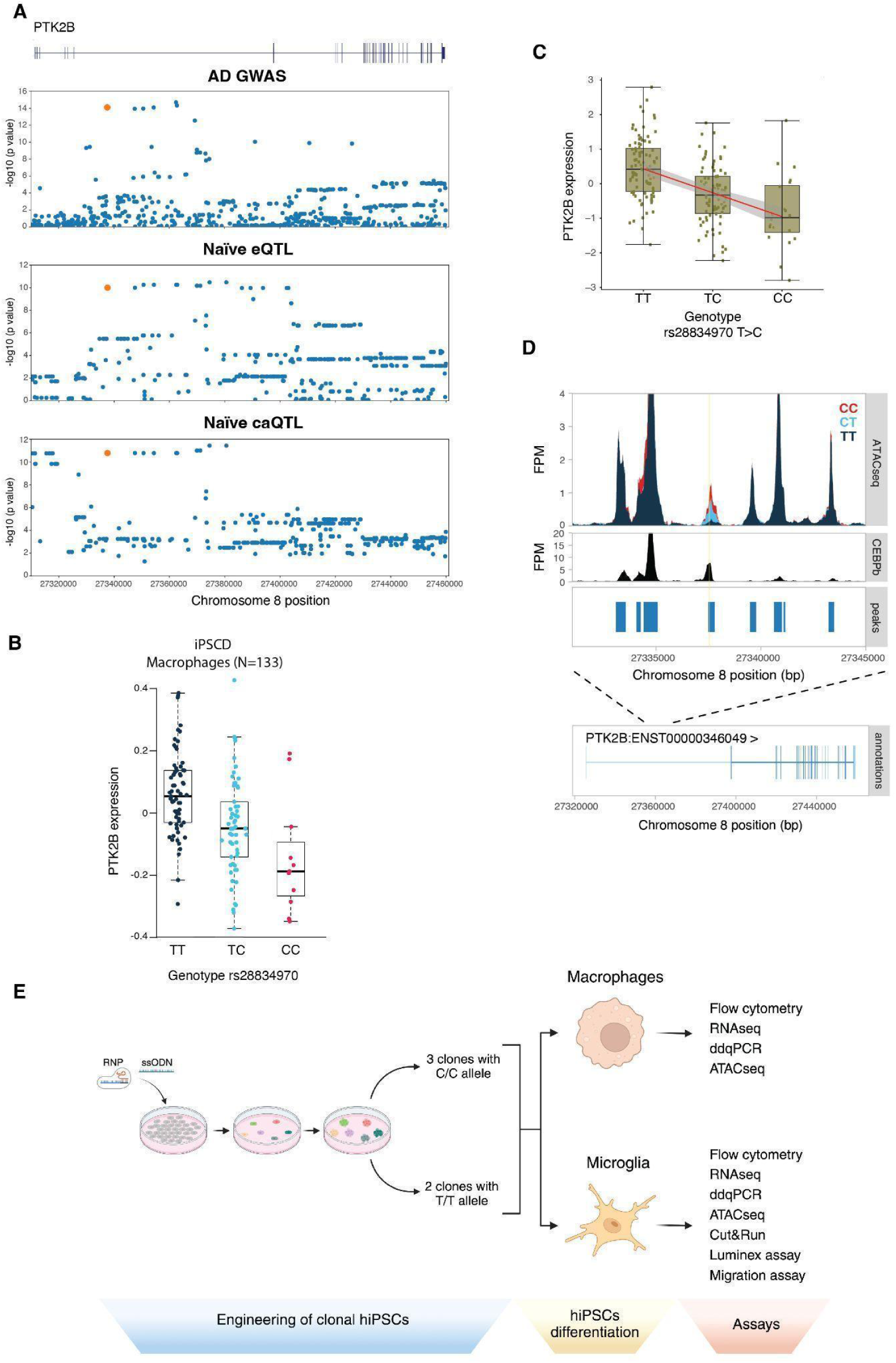
The rs28834970 variant in PTK2B is predicted to modulate an intronic enhancer. A) *PTK2B* gene structure and Manhattan plots for Alzheimer’s Disease (AD) GWAS hits (top panel), eQTL in naïve iPSC-derived macrophages (second panel) and caQTL in naïve iPSC-derived macrophages (bottom panel). P-values replotted from[90]. B) Boxplots showing the expression of the *PTK2B* gene stratified by the rs28834970 genotype in iPSC-derived macrophages. Data replotted from[25]. The y-axis shows normalised expression levels (log TPM value) and each dot shows the expression level of a single sample. C) Boxplots showing the expression of the *PTK2B* gene stratified by the rs28834970 genotype in naïve iPSC-derived macrophages. Data replotted from[91]. The y-axis shows normalised expression levels (log TPM value) and each dot shows the expression level of a single sample. D) ATAC-seq fragment coverage in iPSC-derived macrophages stratified by the rs28834970 genotype (top panel) and CEBPβ ChIP-seq fragment coverage and peaks in primary human macrophages (middle panel). Data replotted from[90]. The rs28834970 variant is highlighted in yellow. The bottom panel shows the structure of the *PTK2B* gene. E) Overview of the experimental design. Created with Biorender.com.

There is some inconsistency in published literature on the directionality of these effects[25,90,91], but this data has been reanalysed in conjunction with the authors, and in additional hiPSC-derived macrophage datasets so we are confident of the directionality of these changes described here. This has confirmed that the C allele of rs28834970 is associated with decreased expression of *PTK2B* in hiPSC-derived macrophages (Fig 1B and Fig 1C), whereas in monocytes and one study of primary microglia, the C allele of rs28834970 is associated with increased expression of *PTK2B* (Supp Fig 1A)[90]. In a recent study this variant was also found to be associated with an increase in PTK2B expression in blood of AD patients and not in any brain region examined[92]. This evidence suggests that the rs28834970 variant is likely causal but highlights the need for further investigation of the role of this variant in AD pathogenesis.

### Characterisation of rs28834970 in hiPSC-derived macrophages and microglia

Given that the PTK2B GWAS signal colocalises with an eQTL/caQTL in hiPSC-derived macrophages, the rs28834970 variant is within a macrophage- and microglia-specific accessible chromatin peak and that PTK2B plays an important role in myeloid cells we decided to focus our investigation on the role of this variant in macrophages and microglia, although we cannot exclude a role in other cell types. We edited the C allele of rs28834970 into a well characterised hiPSC line (KOLF2_C1) that was homozygous for the T allele at this position (and thus the non-risk haplotype) using CRISPR-mediated homology directed repair[93] (Suppl Fig 1B and Methods). This allows us to unambiguously study the role of rs28834970 independently of the other variants in linkage disequilibrium. We subsequently analysed three independent homozygous (C/C) clones and two unedited (T/T) clones, one of which had been through the editing process, and the second being the parental line. These were differentiated in three biological replicates into either macrophages[94,95] or microglia[96,97] followed by a number of assays including transcriptomics (RNAseq), chromatin accessibility (ATACseq) and quantitative digital RT-PCR (dd-qRT-PCR) to measure gene expression (Fig 1E). Differentiation was highly reproducible between the two different genotypes by FACS for macrophage markers CD14, CD16 and CD206 (Supp Fig 2A) and microglia markers CD11b and CD45 (Supp Fig 2B). This was further validated by analysis of macrophage and microglial marker gene expression [96,98] which were mostly consistent with the differentiation trajectory (Supp Fig 2C), including the homeostatic microglial marker P2RY12 that distinguishes between microglia and macrophages and microglial enriched genes e.g. TREM2, GPR34 and C1Qa [96,98]. Moreover, there did not appear to be genotype-dependent changes in expression of canonical markers of macrophage or microglial activation (Supp Fig 2D).

### rs28834970 introduces a novel CEBPB binding site that increases chromatin accessibility at an intronic enhancer in hiPSC-derived macrophages and microglia

We analysed the effect of the rs28834970 allele on chromatin accessibility and function of the intronic enhancer element. ATACseq analysis showed that the C allele created a novel region of accessible chromatin in both macrophages and microglia (Fig 2A), which was consistent with the expectations from the caQTL analysis in hiPSC-derived macrophages (Fig 1D). Analysis of chromatin accessibility within a 600 kb window around the SNP (Supp Fig 3A) showed that there was no change in most peaks, and the only statistically significant change consistent between macrophages and microglia was the new peak over rs28834970 (macrophages: p=1.65e-16 log2FC=2.71, microglia: p=4.19e-12 log2FC=2.77) (Supp Fig 3B). Peaks within the CHRNA2 and CLU genes were significantly reduced in microglia, and an intergenic peak was increased in macrophages, but these were not consistent between the two cell types, or with the expression changes of these genes (Supp Fig 3B and Supplementary Table 1).

**Figure 2.**
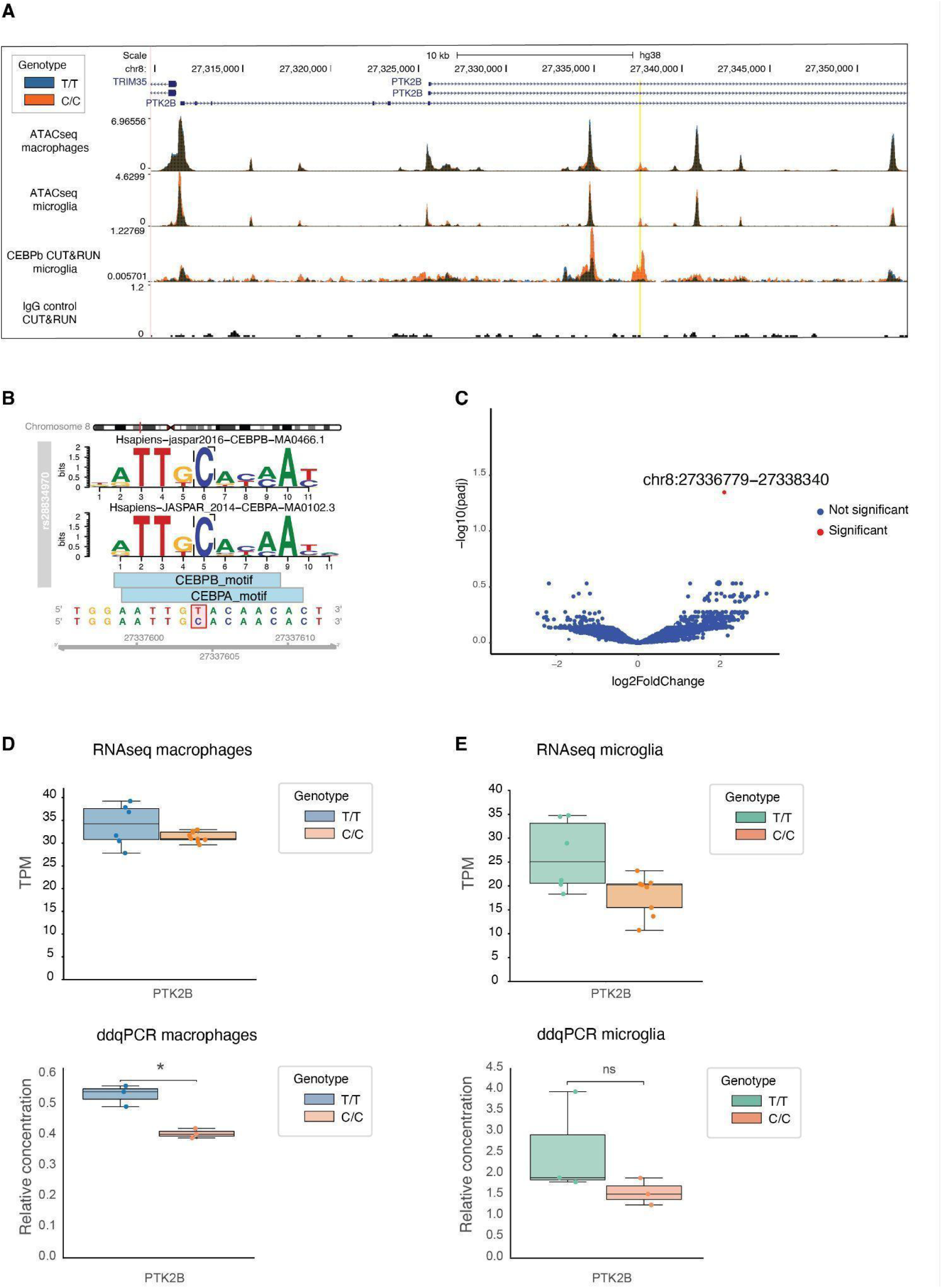
Isogenic edited cell lines show that rs28834970 affects chromatin accessibility and CEBPβ binding at the intronic enhancer in iPSC-derived macrophages and microglia but only modestly decreases *PTK2B* expression. A) Coverage tracks of cells with T/T (orange) and C/C (blue) allele at rs28834970. The tracks are: ATAC-seq in macrophages (upper track), ATAC-seq in microglia (second track), CUT&RUN for CEBPβ in microglia (third track) and IgG negative control for CUT&RUN (bottom track). The rs28834970 variant in *PTK2B* intron is highlighted in yellow. B) Effect of rs28834970 SNP on transcription factor binding probability. Position probability matrices are shown for CEBPβ (top) and CEBPα (middle) as well as the position of the binding motifs on the *PTK2B* sequence (bottom). The plot was generated using motifbreakerR[117]. C) Volcano plot showing fold changes of CEBPβ binding between microglia harbouring the C/C and the T/T allele for all detectable peaks, measured by CUT&RUN. Blue dots indicate peaks with no significant change while red dots represent significantly increased peaks (log2FC>1 and p<0.05). The only significantly increased CEBPβ peak found spans the rs28834970 variant in the *PTK2B* intron. D) Boxplots showing *PTK2B* expression in macrophages harbouring the T/T or C/C allele at rs28834970 measured by RNA-seq (top) and droplet digital q-PCR (ddqPCR) (bottom). Top plot: TPM= transcripts per million. Bottom plot: Relative concentration = copies/ul relative to housekeeping gene (SDHA), n=3 and **p<0.01, unpaired t-test. E) Boxplots showing *PTK2B* expression in microglia harbouring the T/T or C/C allele at rs28834970 measured by RNA-seq (top) and droplet digital q-PCR (ddqPCR) (bottom). Top plot: TPM= transcripts per million. Bottom plot: Relative concentration = copies/ul relative to housekeeping gene (TBP), n=3 and **p<0.01, unpaired t-test. ns= non-significant.

Further analysis of the sequence around rs28834970 showed that the C allele increased the probability of binding of transcription factors CEBPA and CEBPB (Fig 2B). We thus analysed CEBPB binding to DNA by CUT&RUN, which showed an increased binding for the C allele of rs28834970 (p=0.04 log2FC=2.09)(Fig 2A), consistent with CEBPB binding being responsible for the change in chromatin accessibility. Indeed, in a genome-wide analysis, this was the only region that showed a significant change in CEBPb binding between the C/C and T/T alleles at rs28834970 (Fig 2C)(Supplementary Table 2).

We next analysed the effect of rs28834970 on expression of *PTK2B* and the other genes within a 600kb window around this variant (Supp Fig 3A). CHRNA2 and EPHX2 are not expressed in macrophages or microglia and were therefore not investigated. Both RNAseq and digital droplet qPCR (dd-qPCR) analysis showed that expression of *PTK2B* was only modestly reduced by introduction of the C allele in macrophages (RNAseq: p=0.019 and log2FC=-0.2, ddqPCR: p=0.02 and log2FC=-0.39) (Fig 2D) and microglia (RNAseq: p=0.06 log2FC=-0.57, ddqPCR: p=0.29 and log2FC=-0.71) (Fig 2E) and it was only borderline significant with ddqPCR in macrophages and not microglia. However, this effect is consistent in direction between RNAseq and ddqPCR and across microglia and macrophages, and it is in line with the effect of the eQTL (Fig 1B, Fig 1C). The effect on PTK2B expression was the most consistent change in microglia and macrophages within the window around rs28834970, but there was additionally a borderline significant upregulation of CLU expression specifically in macrophages (RNAseq: log2FC=0.72 p=0.03) which was not observed in microglia (Supp Fig 3C). This suggests that there may be an effect of the rs28834970 enhancer on PTK2B and CLU expression, but it is not consistently significant between experiments. This could be due to the small effect size of such common regulatory variants, which makes it difficult to detect in these assays, or to other mutations in the same haplotype contributing further to expression changes in this locus. We also analysed changes in splicing of the PTK2B gene and did not detect any significant changes in splicing of the exons near to the variant (Supp Fig 3D).

Taken together, these results validate that rs28834970 within the PTK2B locus is a regulatory variant that acts through introducing a novel CEBPB binding site that increases chromatin accessibility at the intronic enhancer. There is a subtle decrease in *PTK2B* expression and increase in CLU expression caused by rs28834970 but this is overall not significant so it is difficult to be sure whether the effect of rs28834970 is mediated through PTK2B or CLU expression.

### rs28834970 causes strong effects on the transcriptome including chemokine expression

We next analysed the effects of editing rs28834970 on genome-wide gene expression in macrophages and microglia. In both cell types, principal component analysis showed that although there was a strong clone and batch effect, the two genotypes were well separated by the second principal component (Fig 3A, Fig 3B). Similar results were seen with analysis of global chromatin accessibility, where genotypes were separated by the second principal component (Supp Fig 4A). Analysis of the differentially expressed genes between genotypes showed 690 downregulated and 428 upregulated genes in macrophages (Fig 3A), and 760 downregulated and 328 upregulated in microglia (Fig 3B) (Supplementary Table 3). Importantly, there was a significant (p<0.0001) overlap in the differentially expressed genes between the two cell types (Supp Fig 4B), further corroborating that we were identifying true effects of the genotype, not simply variation between edited clones or differentiation artefacts.

**Figure 3.**
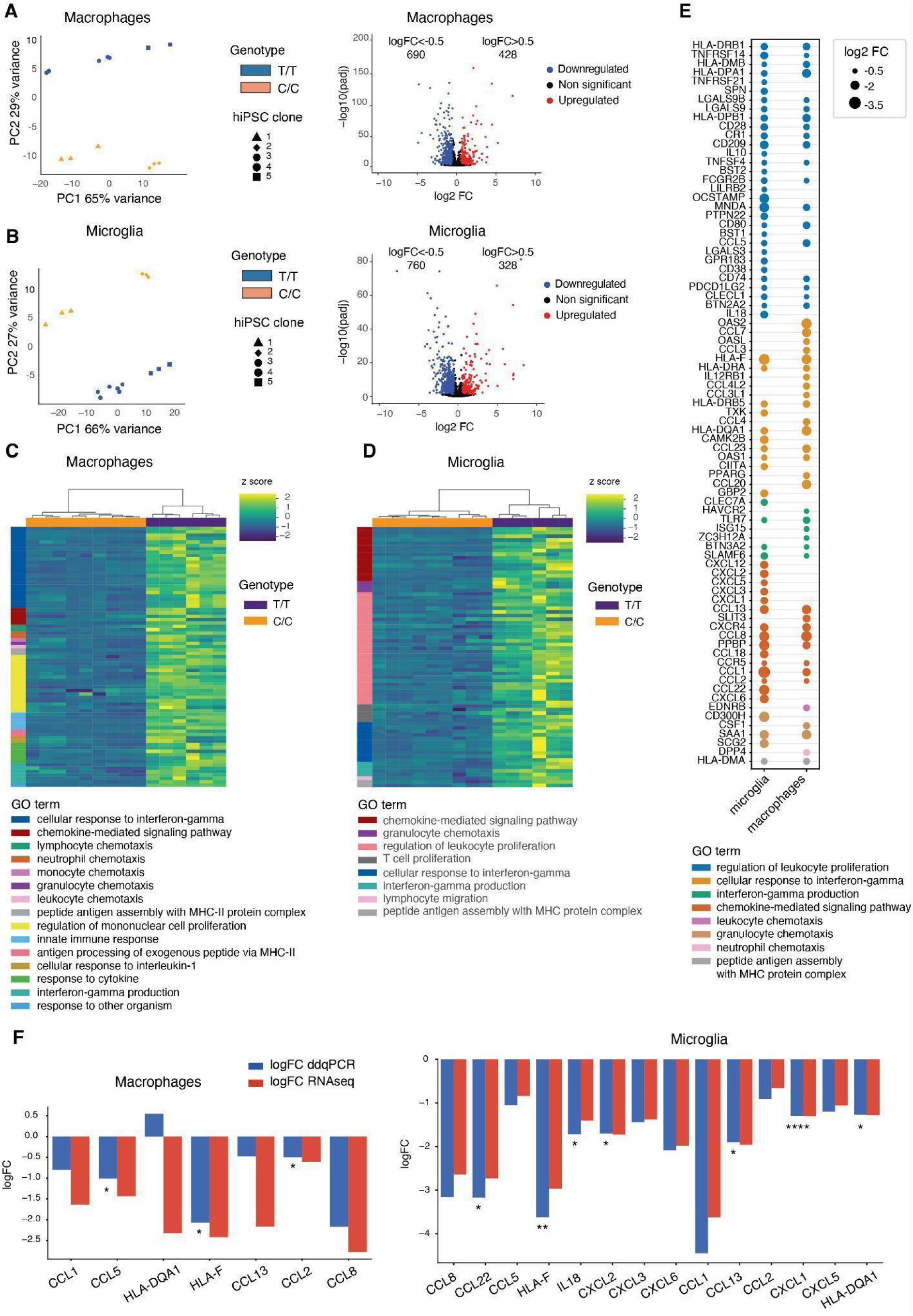
The rs28834970 variant affects global gene expression in iPSC-derived macrophages and microglia. A) Results of differential expression analysis of macrophages harbouring the C/C allele versus the corresponding iPSC-derived cells harbouring the T/T allele at rs28834970. Left panels show Principal component analysis (PCA) of transcriptomes of cells differentiated from two hiPSC clones with the T/T allele and three clones with the C/C allele. Right panels are a volcano plot showing the log2 fold-change (FC) between cells with the C/C and the T/T allele and corresponding p values for all transcribed genes. Significantly upregulated genes (log2FC>0.5 and p<0.05) are shown as red dots, significantly downregulated genes (log2FC<-0.5 and p<0.05) are shown as blue dots and non-significant genes as black dots. B) Results of differential expression analysis of microglia harbouring the C/C allele versus the corresponding iPSC-derived cells harbouring the T/T allele at rs28834970. Left panels show Principal component analysis (PCA) of transcriptomes of cells differentiated from two hiPSC clones with the T/T allele and three clones with the C/C allele. Right panels are a volcano plot showing the log2 fold-change (FC) between cells with the C/C and the T/T allele and corresponding p values for all transcribed genes. Significantly upregulated genes (log2FC>0.5 and p<0.05) are shown as red dots, significantly downregulated genes (log2FC<-0.5 and p<0.05) are shown as blue dots and non-significant genes as black dots. C) Heatmaps showing relative expression (z-scores) of DE genes found in gene ontology (GO) analysis in macrophages with the T/T (orange) and the C/C (purple) allele at rs28834970. GO pathways identified are also annotated. D) Heatmaps showing relative expression (z-scores) of DE genes found in gene ontology (GO) analysis in microglia with the T/T (orange) and the C/C (purple) allele at rs28834970. GO pathways identified are also annotated. E) Dot plot showing log2FC of downregulated genes belonging to the top eight GO pathways in macrophages and microglia. F) Bar graph showing log2FC of downregulated chemokines and HLA genes in macrophages (left) and microglia (right) measured by RNAseq and ddqPCR. ddqPCR: n=3 *p<0.05, **p<0.01, ***p<0.001, ****p<0.0001 unpaired t-test.

We performed gene set enrichment analysis of the differentially expressed genes relative to all genes expressed in the respective cell type. This showed that a number of gene ontology pathways relevant to microglial or macrophage function were enriched in the downregulated gene set including chemokine-mediated signalling, cellular response to interferon gamma (IFNγ) and migration and chemotaxis (Fig 3C, Fig 3D). These pathways were frequently overlapping between microglia and macrophages (Supp Fig 4C), again supporting that these changes were genotype-dependent. Conversely, there were no obviously relevant gene ontology (GO) enrichments in the upregulated gene set (Supp Fig 4D) which may be due to the smaller number of genes analysed.

Further analysis of the downregulated genes confirmed that there was considerable overlap between microglia and macrophages across the different GO categories (Fig 3E). Of note, we found that the Macrophage Receptor With Collagenous Structure (*MARCO*) gene was among the top 3 downregulated genes in both microglia and macrophages with the C/C allele (Supp Fig 4E). MARCO is a putative Aβ receptor that triggers uptake and downstream signalling in microglia [99,100]. Given their role in microglial and macrophage function, we focused on the downregulation of chemokines, MHC complex proteins and surface receptors and validated their expression by dd-qPCR (Supplementary Table 4). All of the chosen genes showed some level of downregulation as expected in microglia, and all except one (HLA-DQA1) were downregulated in macrophages (Fig 3F), with effect sizes generally consistent between dd-qPCR and RNAseq.

### rs28834970 causes changes in chemokine release and migration

Guided by the consistent effects on chemokine and migration-related gene expression in both macrophages and microglia, we next analysed the release of 13 chemokines using a multiplexed assay on microglial cell supernatants from all differentiated clones. In unstimulated cells, we saw significant reduction in production of 11 of the 13 chemokines tested with the C/C allele at rs28834970, consistent with the results of the transcriptomic analysis (Fig 4A). CCL22 was unexpectedly upregulated, and CXCL2 and CXCL5 were unchanged (Fig 4A), which may be a result of the different readouts measured by the two assays (mRNA vs secreted protein).

**Figure 4.**
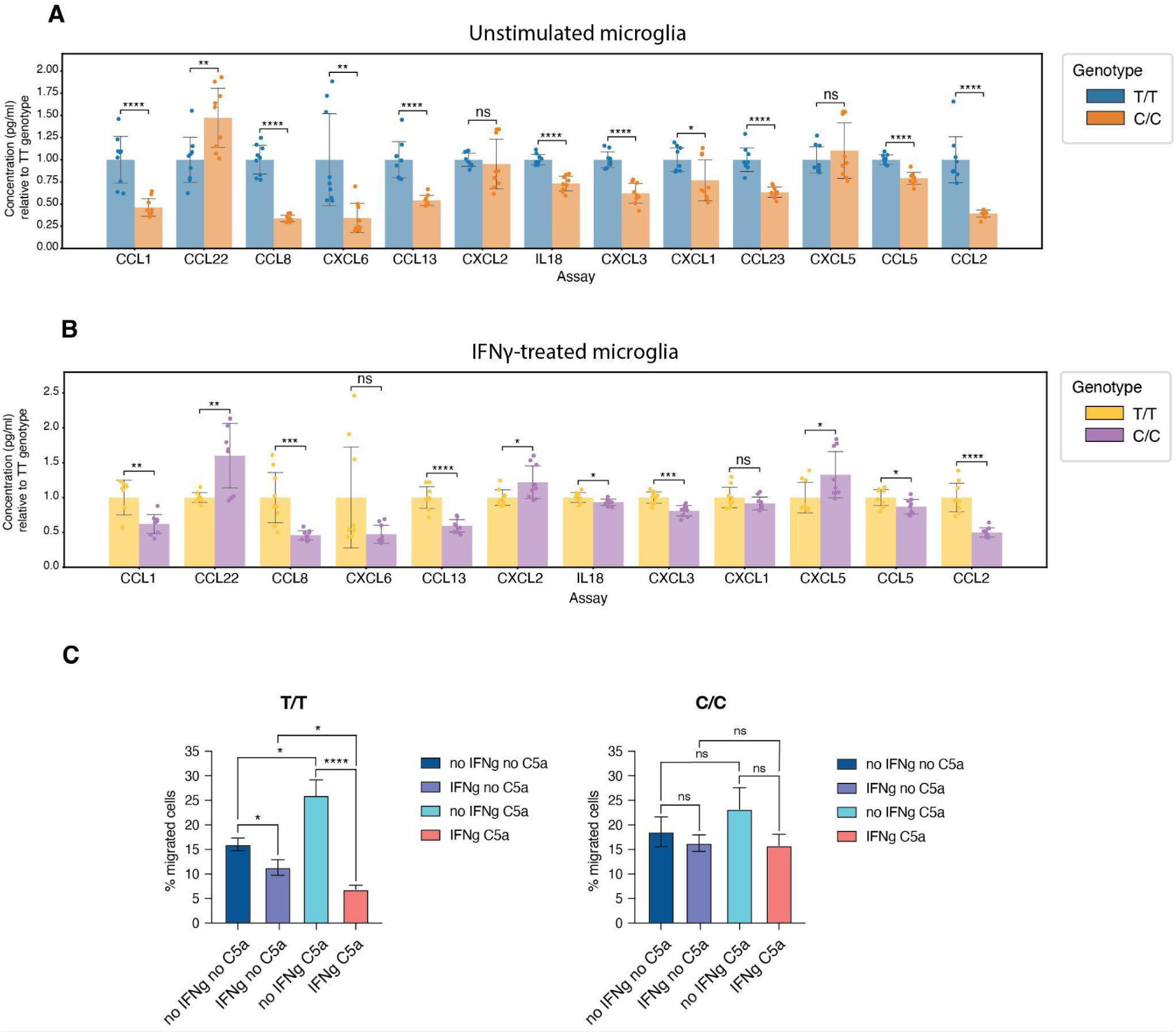
The rs28834970 variant causes changes in chemokine production and migration in iPSC-derived microglia. A) Concentration of downregulated chemokines in unstimulated microglia with the C/C allele at rs28834970 relative to the concentration in cells with the T/T allele, measured by Luminex assay. Results are shown as the mean ± SEM of independent biological replicates (n=3) of each of the five differentiated iPSC clones. ns= non-significant, * p<0.05, **p<0.01, ***p<0.001, ****p<0.0001, an unpaired t-test. B) Concentration of downregulated chemokines in microglia with the C/C allele at rs28834970 relative to the concentration in cells with the T/T allele, stimulated with IFNγ. Results are shown as the mean ± SEM of independent biological replicates (n=3) of each of the five differentiated iPSC clones.. ns= non-significant, * p<0.05, **p<0.01, ***p<0.001, ****p<0.0001, an unpaired t-test. C) Bar graph showing the percentage of migrated microglia harbouring the T/T allele (left) or the C/C allele (right) either unstimulated or stimulated with IFNγ, and with or without the chemoattractant C5a. Results are shown as the mean ± SEM of independent biological replicates (n=3) of each of the five differentiated iPSC clones. 3 technical replicates of each were measured. ns= non-significant, * p<0.05, **p<0.01, ***p<0.001, unpaired t-test.

Next we aimed to evaluate production of these chemokines in stimulated microglia. We treated microglia with IFNγ and lipopolysaccharide (LPS) and analysed whether chemokine production was altered in stimulated microglia. As expected, both stimulations increased production of most (10 of 12) of the chemokines (Supp Fig 5A and Supp Fig 5B). CCL1 production was increased by LPS stimulation in microglia harbouring the C/C allele but not in microglia with the T/T allele (Supp Fig 5B). Similar to what was observed in unstimulated cells, production of most of the chemokines (10 of 12) was reduced in the C/C allele compared to the T/T allele at rs28834970 when microglia were stimulated with IFNγ, but the magnitude of the reduction was reduced (Fig 4B). A similar response was seen upon stimulation with LPS (Supp Fig 5C). Taken together, these data suggest that the C/C allele results in a lower basal chemokine release, and in the presence of stimuli such as IFNγ, this reduction becomes less marked (Supplementary Table 5).

We further investigated the effect of rs28834970 on migration and chemotaxis of microglia differentiated from all iPSC clones, using a transwell assay in the presence or absence of IFNγ and with or without a chemoattractant relevant to microglia, complement 5a (C5a). Microglia harbouring the unedited (T/T) allele showed chemotaxis towards C5a that was inhibited by IFNγ treatment (Fig 4C). Microglia harbouring the C/C alleles at rs28834970 showed no significant chemotaxis towards C5a or inhibition of migration following IFNγ treatment (Fig 4C). Taken together, these data suggest that microglia harbouring the C/C allele are less sensitive to the chemoattractant C5a and to the inhibitory effect of IFNγ on migration.

In summary, we show that the C allele of rs28834970 lowers chemokine release by microglia, reduces chemotaxis towards C5a and alters their response to IFNγ stimulation, perhaps by altering microglia reactivity.

## Discussion

One of the major challenges in interpreting GWAS is to understand the genes and causal variants underlying the genetic association with disease. The use of isogenic cell lines generated by CRISPR in model systems such as hiPSC differentiated cell types offer one way to dissect the role of individual variants in cellular phenotypes independently of other variants in linkage disequilibrium. However, common regulatory variants that are identified in GWAS have rarely been linked to changes in cellular ‘omics or phenotypes. Here we identify a likely causal variant from comparison of Alzheimer’s disease GWAS, eQTL and caQTL studies and engineer this using CRISPR/Cas9 into hiPSCs to generate isogenic cell lines differing in this single variant. We show that the C/C allele of rs28834970 introduces a novel CEBPB binding site that increases chromatin accessibility at an intronic enhancer, consistent with the results of the caQTL analysis in hiPSC-derived macrophages.

Previous eQTL analysis showed that this variant causes a decrease in PTK2B expression in monocytes[94] and macrophages[25,82,83] (Fig 1B and C). However, in our study we observed only a slight downregulation of *PTK2B* in microglia and macrophages expressing the C/C allele, which is only borderline significant in qPCR in macrophages, and doesn’t quite reach significance in microglia or using RNAseq. We also found a subtle upregulation of CLU expression at borderline significance in macrophages but not in microglia expressing the C/C allele, making it difficult to make a definite conclusion on whether rs28834970 affects expression of either PTK2B or CLU or a combination of the two. This could be due to the small effect size making it challenging to study with a limited number of isogenic cell clones, contribution of other SNPs within the same risk haplotype or compensatory regulatory mechanisms that may buffer expression levels. Nevertheless, we found that the C/C allele at rs28834970 caused a large transcriptional response, including downregulation of chemokines and of genes involved in cellular response to IFNγ. This risk variant also resulted in phenotypic changes, such as reduction in chemokine release and decreased chemotaxis towards C5a. Taken together, this suggests that rs28834970 is a functional regulatory variant associated with AD risk at this locus.

We propose that rs28834970 causes a change in microglia reactivity, resulting in a different response to stimulations such as IFNγ. This may impact the ability of the microglia to respond to aggregated proteins such as Aβ plaques and cellular damage in the brain, therefore increasing the risk of AD. Indeed several pro-inflammatory chemokines, such as CCL2, CCL5 and IL18, which are downregulated in microglia with the C/C allele both in naïve state and when stimulated with IFNγ, are produced by microglia in response to Aβ and are important for Aβ-induced microglia chemotaxis[103–105]. Even though overexpression of these chemokines is characteristic of neuroinflammation, correlated with disease progression and found in late stages of AD, knockout of chemokines, such as CCL2, and chemokine receptors, such as CCR2 and CCR5, in mice is associated with increased Aβ deposition and accumulation[47,50–52,106]. It has also been found that patients carrying CCR5Δ32 mutation, which prevents CCR5 surface expression, develop AD at a younger age[107]. Therefore, we hypothesize that in individuals carrying the C/C (risk) allele of rs28834970, downregulation of these chemokines in macrophages and microglia affects Aβ-induced microglia chemotaxis, leukocyte recruitment and clearance of Aβ, and may increase the risk of developing symptomatic AD. The decrease in expression of the putative Aβ receptor MARCO in the C/C allele may also contribute to this effect. We believe this study demonstrates the power of isogenic cell lines to identify and validate individual variants, including common regulatory SNPs identified from GWAS.

## Methods

### Human induced pluripotent stem cells (hiPSC) culture

KOLF2_C1 (https://hpscreg.eu/cell-line/WTSIi018-B-1) is a male hiPSC line generated as part of the HipSci project (Cambridgeshire 1 NRES REC Reference 09/H0304/77, HMDMC 14/013) that has been subcloned from a single cell to ensure homogeneity of the population. KOLF2_C1 cells were cultured in TeSR E8 medium (StemCell Technologies) on Synthemax (Corning) (final amount 2 μg/cm^2^) and routinely passaged 1:10 every 5 days using Gentle Cell Dissociation Reagent (StemCell Technologies).

### Engineering rs28834970 C/C allele in hiPSCs

The C/C risk allele at variant rs28834970 was engineered in hiPSCs by nucleofection of ribonucleoprotein (RNP) complex containing full-length chemically modified synthetic guide RNA and eSpCas9, along with a ssODN repair template, as previously described[93].

Briefly, eSpCas9[108] was expressed and purified from Escherichia coli using a His-tag. KOLF2_C1 cells were cultured ahead of nucleofection in TeSR E8 medium (StemCell Technologies) on Synthemax (Corning) (final amount 5 μg/cm^2^). eSpCas9 (5μl, 20 μg) was mixed with full-length chemically modified guide RNA (AGTGGAATTGTACAACACTG) (Synthego) (5 μl, 225 pmol) at room temperature for 20 min for RNP complexes to form, followed by addition of the ssODN repair template (CTGGCAAGACTAATCTACTTTCTATTTTTATGGATCTGCCTTTTCTGGTCATTCCATATAAGTGGAATTGCACAACACTGTGGCCTTTCGCGACGGCTGCTTTCACTTAGCACAATGTTTTGAAACTTCCTCCATGTTGT) (5 μl, 500 pmol) just before the nucleofection. Cells were detached using Accutase (StemCell Technologies) and dissociated into a single cell suspension. 1 × 10^6^ cells were resuspended in P3 buffer (Lonza), mixed with the RNP/template complex and nucleofected using Lonza 4D-Nucleofector with program CA137. After nucleofection cells were plated onto a 10 cm dish coated with Synthemax (5 μg/cm^2^) with TeSR-E8 supplemented with CloneR (StemCell Technologies). Cells were cultured to 80% confluency, detached using Accutase and dissociated into single cells for subcloning. 5000 cells were plated in a 10 cm dish and expanded, and colonies were picked for genotyping and freezing. Colonies were screened for introduction of the homozygous C allele by high throughput sequencing of an amplicon spanning the rs28834970 variant using an Illumina MiSeq instrument. Final cell lines were validated using Sanger sequencing. Clones used in downstream assays include: three homozygous (C/C) clones (named C8, C11 and D3), one unedited (T/T) picked clone (B11) and the parental KOLF2_C1 line.

### hiPSC differentiation to macrophages

iPSCs were differentiated to macrophages as previously described[94,95]. Briefly, KOLF2_C1 cells were cultured in Essential E8 medium (Gibco) on Vitronectin (Gibco, 100x) and split onto a feeder layer of Mouse Embryonic Fibroblasts (MEFs) (Merck) using Gentle Cell Dissociation Reagent (StemCell Technologies). hiPSCs were cultured on the feeder layer in hiPSC base medium, containing Advanced DMEM F12 (Gibco), KnockOut Serum Replacement (Gibco), 2mM Glutamax (Gibco), 100 U/ml penicillin/streptomycin (Gibco), 0.055 mM 2-Mercaptoethanol (Sigma) and 4ng/ml of Recombinant Human FGF (R&D Systems), until the colonies were sufficiently large. HiPSC colonies were lifted using a solution of equal amounts of Collagenase type IV (ThermoFisher Scientific) and Dispase type II (ThermoFisher Scientific) for 15 minutes at 37°C. Detached colonies were plated in 10 cm low adherent dishes in hiPSC base medium without FGF to form Embryoid Bodies (EBs). After five days, EBs were harvested and plated on gelatin coated 10 cm dishes in EB medium, containing X-Vivo 15 medium (Lonza), 2 mM Glutamax (Gibco), 100 U/ml penicillin/streptomycin (Gibco), 0.055mM β-mercaptoethanol (Sigma), 50 ng/ml rhM-CSF (R&D Systems) and 25 ng/ml rhIL-3 (R&D Systems). Macrophage precursor cells could be routinely harvested from dishes containing EBs every 6/7 days for up to 3/4 months. Precursors were differentiated for 7 days by plating 15,000 cells/cm^2^ in macrophage media, containing RPMI 1640 medium (Gibco), 10% FBS (Thermo Fisher Scientific), 2 mM Glutamax (Gibco), 100 U/ml penicillin/streptomycin (Gibco) and 100 ng/ml rhM-CSF (R&D Systems).

### HiPSCs differentiation to microglia

HiPSCs were differentiated to microglia as previously described[96]. Briefly, KOLF2_C1 cells were cultured in Essential E8 medium (Gibco) on Synthemax (Corning) (final amount 5 μg/cm^2^). Two days after passaging, cells were detached using Accutase (StemCell Technologies) and dissociated into a single cell suspension. 10,000 hiPSCs were plated into each well of a 96-well plate in EB medium, containing Essential E8 medium (Gibco), 10 uM Y-27632 (Dihydrochloride)(StemCell technologies), 50 ng/ml rhBMP-4 (Perprotech), 20 ng/ml rhSCF (Perprotech) and 50 ng/ml rhVEGF (Perprotech). The 96-well plate was span down at 300xg for 3 minutes and incubated for 4/5 days for EB formation. EBs were then transferred to tissue culture dishes coated in 0.1% gelatin (Sigma) and cultured in Microglia Precursors media, containing X-Vivo 15 medium (Lonza), 2 mM Glutamax (Gibco), 100 U/ml penicillin/streptomycin (Gibco), 0.055 mM β-mercaptoethanol (Sigma), 100 ng/ml rhM-CSF (Perprotech) and 25 ng/ml rhIL-3 (Cell Guidance Systems). Microglia precursor cells could be routinely harvested from dishes containing EBs every 6/7 days for up to 3 months. Precursors were differentiated for 7 days by plating at a density of 180,000 cells/cm^2^ in microglia media, containing RPMI 1640 medium (Gibco), 10% FBS (Thermo Fisher Scientific), 2 mM Glutamax (Gibco), 100 U/ml penicillin/streptomycin (Gibco), 10 ng/ml rhGM-CSF (Perprotech) and 100 ng/ml rhIL-34 (Perprotech).

### Fluorescence activated cell sorting analysis

Microglia or macrophages were detached using Accutase (StemCell Technologies) and washed twice in PBS. 1×10^5^ cells were incubated with Human TruStain FcX (Fc Receptor Blocking Solution) (Biolegend) for 30 minutes at 4°C in the dark. Microglia were stained with 5 ul each of PE-anti human CD45 antibody clone HI30 (Biolegend) and APC-anti human CD11b antibody clone ICRF44 (Biolegend) for 30 minutes at 4°C. Macrophages were instead stained with 5 ul of either APC/Cy7-anti human CD16 antibody clone 3G8 (Biolegend), APC-anti human CD206 antibody (BD Biosciences) or PE-anti human CD14 antibody (BD Biosciences). Isotype controls were stained with either 0.2 mg/ml APC-mouse IgG1 k(BD Biosciences), 0.2 mg/ml of PE-mouse IgG2A (BD Biosciences) or 0.2 mg/ml APC/Cy7 mouse IgG1 k (BD Biosciences). Cells were then washed three times in FACS buffer (5% FBS in D-PBS) and resuspended in 200 *μ*l of DAPI solution (1 ug/ml in PBS) and analysed. All samples were analysed using a BD LSRll instrument. An unpaired t-test was used to test the difference between samples from cells with the T/T and the C/C allele at rs28834970.

### RNA extraction

TRI Reagent (Zymo) was added direct on cell layers previously washed in PBS and mixed. RNA was extracted using the Direct-zol RNA Miniprep kit (Zymo) following the manufacturer’s protocol. The optional in-column DNase digest was performed, and RNA was eluted in 50 μl DNAse/RNAse-free water. We made cDNA from 1 μg RNA using Superscript IV (Thermo-Fisher) according to the manufacturer’s protocol.

### Reverse transcription

1 μg RNA was reverse transcribed using Superscript IV (Thermo-Fisher) and random hexamers, according to the manufacturer’s protocol.

### RNA sequencing of hiPSC-derived macrophages and microglia

Transcriptome libraries for hiPSC-derived macrophages and microglia were generated with the Illumina TruSeq stranded RNAseq kit (polyA) and all samples were sequenced using Illumina HiSeq with 50 million mapped reads on average for each sample.

Reads were mapped to the human genome (GRCh38/hg38) using the BWA software[109] and coverage tracks were visualised on the UCSC genome browser using bigwig files, made with bamCoverage[110] and normalised by Counts per Million (CPM). The number of reads per transcript were counted using the featureCounts software[111] and a custom R script was used to calculate Transcripts per Million (TPM). For RNAseq analysis of macrophages, TPMs were corrected for batch effects using limma function limma::removeBatchEffect[112]. For microglia, all samples were differentiated and sequenced in the same batch. Raw read counts per transcript were used to analyse differential gene expression between cells with the T/T and the C/C allele at rs28834970, using the package DESeq2[113]. DESeq2 was also used to generate PCA plots of variance stabilizing transformation (vst) transformed counts. To investigate whether the rs28834970 variant had an effect on PTK2B splicing, we used EdgeR[114] to calculate differential exon usage between microglia harbouring the C/C and the T/T allele. Gene ontology (GO) analysis of differential expressed genes was carried out using g:Profiler package in R[115].

### ATACseq of hiPSC-derived macrophages and microglia

hiPSC-derived macrophages and microglia were detached using a solution of 4 mg/ml Lidocaine hydrochloride monohydrate (Sigma) and 5mM EDTA (ThermoFisher scientific) for 15 minutes at 37°C. ATACseq was performed as previously described[116]. Briefly cells were lysed using sucrose buffer containing 10 mM Tris-Cl pH 7.5, 3 mM CaCl_2_, 2 mM MgCl_2_, 0.32 M sucrose and permeabilised using 0.5% Triton-X-100 (Sigma). Nuclei were recovered by spinning at 450xg for 5 minutes at 4°C and tagmentation was performed using Illumina Tagment DNA TDE1 Enzyme and Buffer Kits for 30 minutes at 37°C37C. 5 *μ*l of TDE1 enzyme was used per 100,000 cells. MinElute PCR Purification Kit buffer PB (Qiagen) was added to stop the reaction and tagmented DNA was purified and eluted in 10 *μ*l of EB. The whole volume was PCR amplified and dual indexed Illumina adapters were added (primer sequences from[116]). Amplified libraries were gel purified and sequenced using an Illumina Novaseq 6000 with 30 million reads on average per sample.

Reads were mapped to the human genome (GRCh38/hg38) using the BWA software[109] and duplicate reads were removed using picard tools MarkDuplicates. Coverage tracks were visualised on the UCSC genome browser using bigwig files, made with bamCoverage[110] and normalised by Counts per Million (CPM). The number of mapped reads per sample was determined and all files were subsampled to the number of reads of the lowest sample, using samtools view[117]. ATACseq peaks were called using MACS2[118] with flags --shift -100 --extsize 200 for window extension. A merged file of all called peaks in all samples was created and the number of reads under each peak was counted in all samples using bedtools coverageBed function[119]. Genome-wide analysis of differential chromatin accessibility between cells harbouring the C/C and the T/T allele at rs28834970 was performed using DESeq2[113]. DESeq2 was also used to generate PCA plots of variance stabilizing transformation (vst) transformed counts. Peaks were annotated to genes using CHIPseeker functiomannotatePeak[120].

### CUT & RUN assay

Microglia were harvested using a solution of 4mg/ml Lidocaine hydrochloride monohydrate (Sigma) and 5mM EDTA (ThermoFisher Scientific) for 15 minutes at 37°C37C. CUT&RUN for CEBPβ was performed using CUTANA CUT&RUN Kit (EpiCypher) according to the manufacturer’s instructions. 400,000 microglia were used per sample and permeabilization was optimised using Digitonin (EpiCypher) 0.01%. 1ug of C/EBP beta Antibody (H-7) sc-7962 (Santa Cruz Biotechnology) was used per sample and one control sample was treated in parallel with 1 *μ*l of CUTANA Rabbit IgG CUT&RUN Negative Control Antibody. 0.5ng of E. coli Spike-in DNA (EpiCypher) was added to each sample to normalise reads to control for experimental variability and sequencing depth. Around 5 ng of purified CUT&RUN-enriched DNA was used to prepare NGS libraries using the NEBNext Ultra II DNA Library Prep Kit (NEB) with NEBNext Multiplex Oligos for Illumina (Dual Index Primers Set 1) (NEB). All libraries were sequenced 150bp paired end using an Illumina Novaseq 6000 with 15 million reads on average per sample.

Reads were trimmed to exclude Illumina adapter sequences using trim_galore and mapped to the human genome (GRCh38/hg38) using the bowtie2 software[121]. The number of human mapped reads per sample was determined and all files were subsampled to the number of reads of the lowest sample, using samtools view[117]. Trimmed reads were also mapped to the E. coli genome using bowtie2 and reads mapped to the human genome were normalised to spiked-in E.coli DNA using a previously published script (https://github.com/Henikoff/Cut-and-Run/blob/master/spike_in_calibration.csh) integrated into a custom bash script (https://github.com/ericabello/PTK2B_rs28834970/blob/main/Cut_Run_CEBPb_micro/cmds_CutRun_CEBPb_micro.sh). Coverage tracks were visualised on the UCSC genome browser using bigwig files made from E.coli normalised bedgraph files of fragments 1-1000bp. Bigwigs were made using UCSC bedGraphToBigWig. Peaks in all samples containing the CEBPβ antibody were called using the SEACR software[122] and the IgG negative control sample was used as input to identify the threshold value at which the percentage of target versus IgG signal is maximized. A merged file of all CEBPβ peaks in all samples was created and the number of reads under each peak was counted in all samples using bedtools coverageBed function[119]. Genome-wide analysis of differential CEBPβ binding between microglia harbouring the C/C and the T/T allele at rs28834970 was performed using DESeq2[113].

### Digital droplet qPCR

50 ng of cDNA per sample was mixed with 10 μl of ddPCR Supermix for probes (no dUTP) (Biorad), 1 μl of FAM-tagged TaqMan assay for detection of the gene of interest and 1 μl of VIC-tagged Taqman assay for a housekeeping gene control. The housekeeping control gene was chosen to match the level of expression of the gene of interest in each cell type.

Taqman assays list (ThermoFisher Scientific), containing probes and primers:

*Macrophages*

| gene | FAM probe ID | control gene | VIC control probe ID |
| --- | --- | --- | --- |
| CCL1 | Hs00171072_m1 | TBP | Hs00427620_m1 |
| CCL8 | Hs00271615_m1 | SDHA | Hs00188166_m1 |
| CCL13 | Hs00234646_m1 | GAPDH | Hs99999905_m1 |
| CCL5 | Hs99999048_m1 | TBP | Hs00427620_m1 |
| CCL2 | Hs00234140_m1 | UBC | Hs00824723_m1 |
| TRIM35 | Hs01093979_m1 | TBP | Hs00427620_m1 |
| CLU | Hs00156548_m1 | TBP | Hs00427620_m1 |
| SCARA3 | Hs00939871_m1 | TBP | Hs00427620_m1 |
| PTK2B | Hs01559703_m1 | SDHA | Hs00188166_m1 |
| HLA-DQA1 | Hs03007426_mH | TBP | Hs00427620_m1 |
| HLA-F | Hs01587840_m1 | TBP | Hs00427620_m1 |

*Microglia*

| gene | FAM probe ID | control gene | VIC control probe ID |
| --- | --- | --- | --- |
| CCL1 | Hs00171072_m1 | HPRT1 | Hs02800695_m1 |
| CCL22 | Hs00171080_m1 | TBP | Hs00427620_m1 |
| CCL8 | Hs00271615_m1 | TBP | Hs00427620_m1 |
| CXCL6 | Hs00237017_m1 | TBP | Hs00427620_m1 |
| CCL13 | Hs00234646_m1 | UBC | Hs00824723_m1 |
| CXCL2 | Hs00601975_m1 | TBP | Hs00427620_m1 |
| IL18 | Hs01038788_m1 | TBP | Hs00427620_m1 |
| CXCL3 | Hs00171061_m1 | TBP | Hs00427620_m1 |
| CXCL1 | Hs00236937_m1 | SDHA | Hs00188166_m1 |
| CXCL5 | Hs00171085_m1 | SDHA | Hs00188166_m1 |
| CCL5 | Hs99999048_m1 | TBP | Hs00427620_m1 |
| CCL2 | Hs00234140_m1 | GAPDH | Hs99999905_m1 |
| TRIM35 | Hs01093979_m1 | TBP | Hs00427620_m1 |
| CLU | Hs00156548_m1 | TBP | Hs00427620_m1 |
| SCARA3 | Hs00939871_m1 | TBP | Hs00427620_m1 |
| PTK2B | Hs00169444_m1 | TBP | Hs00427620_m1 |
| HLA-DQA1 | Hs03007426_mH | TBP | Hs00427620_m1 |
| HLA-F | Hs01587840_m1 | TBP | Hs00427620_m1 |

Droplets were generated for each reaction using a QX200 Droplet Generator (Biorad) and transferred to 96 well plates. Sealed plates were incubated in a thermocycler and cDNA was PCR amplified with the following steps: 95°C for 10 min then 95°C for 30 sec and 60°C for 1 min for 40 cycles and finally 98°C for 10 min. The resulting FAM and VIC fluorescence was read using a QX200 Droplet Reader (Biorad) and the fraction of positive droplets in the sample was determined. Poisson statistics were used to determine the absolute number of copies per μl in each sample for both target and housekeeping control genes. The number of copies of target gene per sample was normalised by the control gene for each reaction and an unpaired t-test Benjamini-Hochberg multiple testing correction was used to test the difference between samples from cells with the T/T and the C/C allele at rs28834970.

### Microglia stimulation and Luminex assay

hiPSC-derived microglia at day 6 of differentiation were stimulated with IFN*γ* (R&D Systems) 200 ng/ml or Lipopolysaccharide (LPS) (Invitrogen) 100 ng/ml for 24 hours. Culture plates were spun at 4,000 rpm for 20 min at 4°C and media collected. Harvested media was diluted 1:20 for unstimulated controls and 1:250 for stimulated samples. Luminex assay was performed using ProcartaPlex Human Basic Kit (Invitrogen) and a panel of ProcartaPlex Human Simplex (Invitrogen) beads, according to the manufacturer’s recommendations. Plates were read using a Luminex MAGPIX instrument. Media from cells with the T/T and C/C allele were ran on the same plate, allowing direct comparison of each protein concentration across genotypes. An unpaired t-test with Benjamini-Hochberg multiple testing correction was used to test the difference in concentration (pg/ml) between different samples. The propagation of error for log fold change values of chemokine concentration between unstimulated and stimulated microglia was calculated using the Python package uncertainties (https://pythonhosted.org/uncertainties/).

### Microglia stimulation and migration assay

HiPSCs-derived microglia at day 7 of differentiation were stimulated with IFN*γ* (R&D Systems) 100 ng/ml for 72 hours. At day 10, cells were detached using a solution of 4 mg/ml Lidocaine hydrochloride monohydrate (Sigma) and 5 mM EDTA (Gibco) and 5.5 × 10^4^ cells in 100 *μ*l macrophage media were seeded onto each transwell (PET with 5 μm pores, Sarstedt) in a 24-well plate. Cells were incubated for 15 minutes to settle and 600 *μ*l of macrophage media containing 3 nM human recombinant C5a was added beneath the transwells. The plate was incubated for 10 hours to allow cell migration. The transwell was then rinsed in PBS, transferred to a fresh 24-well plate, and fixed with 4% paraformaldehyde for 20 min at RT. Cells on either side of the transwell membrane were stained with NucBlue (Invitrogen) and imaged with an EVOS FL Auto automated microscope (Thermo Fisher) set up to take images of each transwell in full (top image). The transwells were swabbed with a cotton wool bud to remove cells on the top surface, leaving behind only migrated cells, transferred into a fresh plate and imaged again with the same settings (bottom image). Cell counting was performed with CellProfiler 3.0 software[123] and the percentage of migrated cells per transwell was calculated as: (no. cells in bottom image) ÷ (no. cells in top image) × 100. Stimulations were performed in triplicate and the average for each sample across wells was taken. An unpaired t-test was used to test the difference between different samples.

## Data availability

Raw sequencing data is available at ENA https://www.ebi.ac.uk/ena/browser/search with accession number ERP024123 for ATACseq and Cut&Run data and ERP024128 for RNAseq data. A description of the sample IDs can be found in Supplementary Table 6.

All analysis scripts are available at https://github.com/ericabello/PTK2B_rs28834970.

## Acknowledgements

This work was funded by OpenTargets [OTAR037] and Wellcome [206194]. We thank the Sanger Institute flow cytometry core facility for flow cytometry analysis. We acknowledge Scientific Operations at the Sanger Institute for support in next generation sequencing and quality control. We thank members of the Bassett lab and Magdalena Strauss for helpful discussions and support, and Marta Perez-Alcantara for supporting computational analysis.

## Author contributions

A.B. and E.B. conceived the work and designed experiments, K.L. made the hiPSC isogenic cell lines, E.B., S.I., J.S. executed the experiments, E.B., K.A., N.K., J.S., N.I.P. analysed data, E.B. and A.B. wrote the manuscript with input from all other authors.

## Declaration of interests

A.B. is a founder of EnsoCell therapeutics. N.I.P. was an employee of GSK at the time the manuscript was submitted. J.S. was an employee of Illumina at the time the manuscript was submitted.

## Materials and Correspondance

All correspondence and requests for materials should be addressed to Andrew Bassett or Erica Bello.

## Ethics statement

Derivation of the KOLF2 (https://hpscreg.eu/cell-line/WTSIi018-B-1) hiPSC line was as part of the HipSci project that was approved by the NRES research ethics committee (Cambridgeshire 1 NRES REC Reference 09/H0304/77, HMDMC 14/013) and work was performed in accordance with appropriate ethical standards.

## Notes

### Summary of Updates

Revised figures to include individual datapoints, altered discussion of data and included additional raw data in new supplementary tables.

## References

1. Gatz M, Reynolds CA, Fratiglioni L, Johansson B, Mortimer JA, Berg S, et al. Role of genes and environments for explaining Alzheimer disease. Arch Gen Psychiatry. 2006;63. doi:10.1001/archpsyc.63.2.168

2. Knopman DS, Amieva H, Petersen RC, Chételat G, Holtzman DM, Hyman BT, et al. Alzheimer disease. Nature Reviews Disease Primers. 2021;7: 1–21.

3. The complex genetic architecture of Alzheimer’s disease: novel insights and future directions. eBioMedicine. 2023;90: 104511.

4. Bellenguez C, Küçükali F, Jansen IE, Kleineidam L, Moreno-Grau S, Amin N, et al. New insights into the genetic etiology of Alzheimer’s disease and related dementias. Nat Genet. 2022;54. doi:10.1038/s41588-022-01024-z

5. Kunkle BW, Grenier-Boley B, Sims R, Bis JC, Damotte V, Naj AC, et al. Genetic meta-analysis of diagnosed Alzheimer’s disease identifies new risk loci and implicates Aβ, tau, immunity and lipid processing. Nat Genet. 2019;51: 414–430.

6. Schwartzentruber J, Cooper S, Liu JZ, Barrio-Hernandez I, Bello E, Kumasaka N, et al. Genome-wide meta-analysis, fine-mapping and integrative prioritization implicate new Alzheimer’s disease risk genes. Nat Genet. 2021;53. doi:10.1038/s41588-020-00776-w

7. Ghoussaini M, Mountjoy E, Carmona M, Peat G, Schmidt EM, Hercules A, et al. Open Targets Genetics: systematic identification of trait-associated genes using large-scale genetics and functional genomics. Nucleic Acids Res. 2021;49: D1311–D1320.

8. Klein JC, Agarwal V, Inoue F, Keith A, Martin B, Kircher M, et al. A systematic evaluation of the design and context dependencies of massively parallel reporter assays. Nat Methods. 2020;17: 1083–1091.

9. Cooper SE, Schwartzentruber J, Bello E, Coomber EL, Bassett AR. Screening for functional transcriptional and splicing regulatory variants with GenIE. Nucleic Acids Res. 2020;48: e131.

10. Cooper S, Schwartzentruber J, Coomber EL, Wu Q, Bassett A. Screening for functional regulatory variants in open chromatin using GenIE-ATAC. Nucleic Acids Res. 2023. doi:10.1093/nar/gkad332

11. Kwart D, Gregg A, Scheckel C, Murphy EA, Paquet D, Duffield M, et al. A Large Panel of Isogenic APP and PSEN1 Mutant Human iPSC Neurons Reveals Shared Endosomal Abnormalities Mediated by APP β-CTFs, Not Aβ. Neuron. 2019;104. doi:10.1016/j.neuron.2019.07.010

12. Penney J, Ralvenius WT, Tsai L-H. Modeling Alzheimer’s disease with iPSC-derived brain cells. Mol Psychiatry. 2019;25: 148–167.

13. Moore S, Evans LDB, Andersson T, Portelius E, Smith J, Dias TB, et al. APP Metabolism Regulates Tau Proteostasis in Human Cerebral Cortex Neurons. Cell Rep. 2015;11: 689–696.

14. APOE4 Causes Widespread Molecular and Cellular Alterations Associated with Alzheimer’s Disease Phenotypes in Human iPSC-Derived Brain Cell Types. Neuron. 2018;98: 1141–1154.e7.

15. Hall-Roberts H, Agarwal D, Obst J, Smith TB, Monzón-Sandoval J, Di Daniel E, et al. TREM2 Alzheimer’s variant R47H causes similar transcriptional dysregulation to knockout, yet only subtle functional phenotypes in human iPSC-derived macrophages. Alzheimers Res Ther. 2020;12: 151.

16. Soldner F, Stelzer Y, Shivalila CS, Abraham BJ, Latourelle JC, Barrasa MI, et al. Parkinson-associated risk variant in distal enhancer of α-synuclein modulates target gene expression. Nature. 2016;533: 95–99.

17. Volpato V, Smith J, Sandor C, Ried JS, Baud A, Handel A, et al. Reproducibility of Molecular Phenotypes after Long-Term Differentiation to Human iPSC-Derived Neurons: A Multi-Site Omics Study. Stem Cell Reports. 2018;11: 897–911.

18. Labzin LI, Heneka MT, Latz E. Innate Immunity and Neurodegeneration. Annu Rev Med. 2018;69. doi:10.1146/annurev-med-050715-104343

19. Wang H. Microglia Heterogeneity in Alzheimer’s Disease: Insights From Single-Cell Technologies. Front Synaptic Neurosci. 2021;13: 773590.

20. Li Q, Barres BA. Microglia and macrophages in brain homeostasis and disease. Nat Rev Immunol. 2017;18: 225–242.

21. Li Y, Laws SM, Miles LA, Wiley JS, Huang X, Masters CL, et al. Genomics of Alzheimer’s disease implicates the innate and adaptive immune systems. Cell Mol Life Sci. 2021;78: 7397–7426.

22. Nott A, Holtman IR. Genetic insights into immune mechanisms of Alzheimer’s and Parkinson’s disease. Front Immunol. 2023;14: 1168539.

23. Tansey KE, Cameron D, Hill MJ. Genetic risk for Alzheimer’s disease is concentrated in specific macrophage and microglial transcriptional networks. Genome Med. 2018;10. doi:10.1186/s13073-018-0523-8

24. Lopes KP, Snijders GJL, Humphrey J, Allan A, Sneeboer MAM, Navarro E, et al. Genetic analysis of the human microglial transcriptome across brain regions, aging and disease pathologies. Nat Genet. 2022;54. doi:10.1038/s41588-021-00976-y

25. Young AMH, Kumasaka N, Calvert F, Hammond TR, Knights A, Panousis N, et al. A map of transcriptional heterogeneity and regulatory variation in human microglia. Nat Genet. 2021;53. doi:10.1038/s41588-021-00875-2

26. Ulland TK, Colonna M. TREM2 — a key player in microglial biology and Alzheimer disease. Nat Rev Neurol. 2018;14: 667–675.

27. Zhao L. CD33 in Alzheimer’s Disease – Biology, Pathogenesis, and Therapeutics: A Mini-Review. Gerontology. 2018;65: 323–331.

28. Novoa C, Salazar P, Cisternas P, Gherardelli C, Vera-Salazar R, Zolezzi JM, et al. Inflammation context in Alzheimer’s disease, a relationship intricate to define. Biol Res. 2022;55: 1–18.

29. Vogels T, Murgoci A-N, Hromádka T. Intersection of pathological tau and microglia at the synapse. Acta Neuropathologica Communications. 2019;7: 1–25.

30. Rogers J, Strohmeyer R, Kovelowski CJ, Li R. Microglia and inflammatory mechanisms in the clearance of amyloid β peptide. Glia. 2002;40: 260–269.

31. Hansen DV, Hanson JE, Sheng M. Microglia in Alzheimer’s disease. J Cell Biol. 2018;217. doi:10.1083/jcb.201709069

32. Asai H, Ikezu S, Tsunoda S, Medalla M, Luebke J, Haydar T, et al. Depletion of microglia and inhibition of exosome synthesis halt tau propagation. Nat Neurosci. 2015;18. doi:10.1038/nn.4132

33. Hong S, Beja-Glasser VF, Nfonoyim BM, Frouin A, Li S, Ramakrishnan S, et al. Complement and microglia mediate early synapse loss in Alzheimer mouse models. Science. 2016;352. doi:10.1126/science.aad8373

34. Miao J, Ma H, Yang Y, Liao Y, Lin C, Zheng J, et al. Microglia in Alzheimer’s disease: pathogenesis, mechanisms, and therapeutic potentials. Front Aging Neurosci. 2023;15: 1201982.

35. Monoranu CM, Hartmann T, Strobel S, Heinsen H, Riederer P, Distel L, et al. Is There Any Evidence of Monocytes Involvement in Alzheimer’s Disease? A Pilot Study on Human Postmortem Brain. Journal of Alzheimer’s disease reports. 2021;5. doi:10.3233/ADR-210052

36. Malm T, Koistinaho M, Muona A, Magga J, Koistinaho J. The role and therapeutic potential of monocytic cells in Alzheimer’s disease. Glia. 2010;58: 889–900.

37. Gião T, Teixeira T, Almeida MR, Cardoso I. Choroid Plexus in Alzheimer’s Disease-The Current State of Knowledge. Biomedicines. 2022;10. doi:10.3390/biomedicines10020224

38. Park L, Uekawa K, Garcia-Bonilla L, Koizumi K, Murphy M, Pistik R, et al. Brain Perivascular Macrophages Initiate the Neurovascular Dysfunction of Alzheimer Aβ Peptides. Circ Res. 2017;121. doi:10.1161/CIRCRESAHA.117.311054

39. De Schepper S, Ge JZ, Crowley G, Ferreira LSS, Garceau D, Toomey CE, et al. Perivascular cells induce microglial phagocytic states and synaptic engulfment via SPP1 in mouse models of Alzheimer’s disease. Nat Neurosci. 2023;26. doi:10.1038/s41593-023-01257-z

40. Munawara U, Catanzaro M, Xu W, Tan C, Hirokawa K, Bosco N, et al. Hyperactivation of monocytes and macrophages in MCI patients contributes to the progression of Alzheimer’s disease. Immun Ageing. 2021;18: 1–25.

41. Thériault P, ElAli A, Rivest S. The dynamics of monocytes and microglia in Alzheimer’s disease. Alzheimers Res Ther. 2015;7: 41.

42. Yan P, Kim K-W, Xiao Q, Ma X, Czerniewski LR, Liu H, et al. Peripheral monocyte-derived cells counter amyloid plaque pathogenesis in a mouse model of Alzheimer’s disease. J Clin Invest. 2022;132. doi:10.1172/JCI152565

43. Michaud J-P, Bellavance M-A, Préfontaine P, Rivest S. Real-time in vivo imaging reveals the ability of monocytes to clear vascular amyloid beta. Cell Rep. 2013;5: 646–653.

44. Hawkes CA, McLaurin J. Selective targeting of perivascular macrophages for clearance of β-amyloid in cerebral amyloid angiopathy. Proceedings of the National Academy of Sciences. 2009;106: 1261–1266.

45. Fiala M, Cribbs DH, Rosenthal M, Bernard G. Phagocytosis of amyloid-beta and inflammation: two faces of innate immunity in Alzheimer’s disease. J Alzheimers Dis. 2007;11: 457–463.

46. Costarelli L, Malavolta M, Giacconi R, Provinciali M. Dysfunctional macrophages in Alzheimer Disease: another piece of the “macroph-aging” puzzle? Aging. 2017. pp. 1865–1866.

47. Wojcieszak J, Kuczyńska K, Zawilska JB. Role of Chemokines in the Development and Progression of Alzheimer’s Disease. J Mol Neurosci. 2022;72: 1929–1951.

48. Zuena AR, Casolini P, Lattanzi R, Maftei D. Chemokines in Alzheimer’s Disease: New Insights Into Prokineticins, Chemokine-Like Proteins. Front Pharmacol. 2019;10: 622.

49. Azizi G, Khannazer N, Mirshafiey A. The Potential Role of Chemokines in Alzheimer’s Disease Pathogenesis. Am J Alzheimers Dis Other Demen. 2014;29: 415–425.

50. Naert G, Rivest S. CC chemokine receptor 2 deficiency aggravates cognitive impairments and amyloid pathology in a transgenic mouse model of Alzheimer’s disease. J Neurosci. 2011;31: 6208–6220.

51. El Khoury J, Toft M, Hickman SE, Means TK, Terada K, Geula C, et al. Ccr2 deficiency impairs microglial accumulation and accelerates progression of Alzheimer-like disease. Nat Med. 2007;13: 432–438.

52. Mildner A, Schlevogt B, Kierdorf K, Böttcher C, Erny D, Kummer MP, et al. Distinct and non-redundant roles of microglia and myeloid subsets in mouse models of Alzheimer’s disease. J Neurosci. 2011;31: 11159–11171.

53. Xia MQ, Hyman BT. Chemokines/chemokine receptors in the central nervous system and Alzheimer’s disease. J Neurovirol. 1999;5: 32–41.

54. Zhu X, Bao Y, Guo Y, Yang W. Proline-Rich Protein Tyrosine Kinase 2 in Inflammation and Cancer. Cancers . 2018;10. doi:10.3390/cancers10050139

55. Beck TN, Nicolas E, Kopp MC, Golemis EA. Adaptors for disorders of the brain? The cancer signaling proteins NEDD9, CASS4, and PTK2B in Alzheimer’s disease. Oncoscience. 2014;1: 486.

56. de Pins B, Mendes T, Giralt A, Girault J-A. The Non-receptor Tyrosine Kinase Pyk2 in Brain Function and Neurological and Psychiatric Diseases. Front Synaptic Neurosci. 2021;13: 749001.

57. Day RN, Day KH, Pavalko FM. Direct visualization by FRET-FLIM of a putative mechanosome complex involving Src, Pyk2 and MBD2 in living MLO-Y4 cells. PLoS One. 2021;16. doi:10.1371/journal.pone.0261660

58. Girault JA, Costa A, Derkinderen P, Studler JM, Toutant M. FAK and PYK2/CAKbeta in the nervous system: a link between neuronal activity, plasticity and survival? Trends Neurosci. 1999;22. doi:10.1016/s0166-2236(98)01358-7

59. Ivankovic-Dikic I, Grönroos E, Blaukat A, Barth BU, Dikic I. Pyk2 and FAK regulate neurite outgrowth induced by growth factors and integrins. Nat Cell Biol. 2000;2. doi:10.1038/35023515

60. Kong L, Wang B, Yang X, He B, Hao D, Yan L. Integrin-associated molecules and signalling cross talking in osteoclast cytoskeleton regulation. J Cell Mol Med. 2020;24. doi:10.1111/jcmm.15052

61. Miao L, Xin X, Xin H, Shen X, Zhu Y-Z. Hydrogen Sulfide Recruits Macrophage Migration by Integrin β1-Src-FAK/Pyk2-Rac Pathway in Myocardial Infarction. Sci Rep. 2016;6: 1–10.

62. Okigaki M, Davis C, Falasca M, Harroch S, Felsenfeld DP, Sheetz MP, et al. Pyk2 regulates multiple signaling events crucial for macrophage morphology and migration. Proc Natl Acad Sci U S A. 2003;100. doi:10.1073/pnas.1834348100

63. Du Q-S, Ren X-R, Xie Y, Wang Q, Mei L, Xiong W-C. Inhibition of PYK2-induced actin cytoskeleton reorganization, PYK2 autophosphorylation and focal adhesion targeting by FAK. J Cell Sci. 2001;114: 2977–2987.

64. Das P, Pal S, Oldfield CM, Thillai K, Bala S, Carnevale KA, et al. A PKCβ-LYN-PYK2 Signaling Axis Is Critical for MCP-1-Dependent Migration and Adhesion of Monocytes. J Immunol. 2021;206. doi:10.4049/jimmunol.1900706

65. Yamamoto S, Shimizu S, Kiyonaka S, Takahashi N, Wajima T, Hara Y, et al. TRPM2-mediated Ca2+ influx induces chemokine production in monocytes that aggravates inflammatory neutrophil infiltration. Nat Med. 2008;14: 738.

66. Alawieyah SMS, Sim JA, Neubrand VE, Jiang LH. A critical role of TRPM2 channel in Aβ42 -induced microglial activation and generation of tumor necrosis factor-α. Glia. 2018;66. doi:10.1002/glia.23265

67. Lambert J-C, Ibrahim-Verbaas CA, Harold D, Naj AC, Sims R, Bellenguez C, et al. Meta-analysis of 74,046 individuals identifies 11 new susceptibility loci for Alzheimer’s disease. Nat Genet. 2013;45: 1452–1458.

68. Beecham GW, Hamilton K, Naj AC, Martin ER, Huentelman M, Myers AJ, et al. Genome-wide association meta-analysis of neuropathologic features of Alzheimer’s disease and related dementias. PLoS Genet. 2014;10. doi:10.1371/journal.pgen.1004606

69. Jiao B, Liu X, Zhou L, Wang MH, Zhou Y, Xiao T, et al. Polygenic Analysis of Late-Onset Alzheimer’s Disease from Mainland China. PLoS One. 2015;10. doi:10.1371/journal.pone.0144898

70. Li YQ, Tan MS, Wang HF, Tan CC, Zhang W, Zheng ZJ, et al. Common variant in PTK2B is associated with late-onset Alzheimer’s disease: A replication study and meta-analyses. Neurosci Lett. 2016;621. doi:10.1016/j.neulet.2016.04.020

71. Wang X, Lopez O, Sweet RA, Becker JT, DeKosky ST, Barmada MM, et al. Genetic Determinants of Survival in Patientswith Alzheimer’s Disease. J Alzheimers Dis. 2015;45. doi:10.3233/JAD-142442

72. Nettiksimmons J, Tranah G, Evans DS, Yokoyama JS, Yaffe K. Gene-based aggregate SNP associations between candidate AD genes and cognitive decline. Age . 2016;38. doi:10.1007/s11357-016-9885-2

73. Lawingco T, Chaudhury S, Brookes KJ, Guetta-Baranes T, Guerreiro R, Bras J, et al. Genetic variants in glutamate-, Aβ-, and tau-related pathways determine polygenic risk for Alzheimer’s disease. Neurobiol Aging. 2021;101. doi:10.1016/j.neurobiolaging.2020.11.009

74. Rajabli F, Benchek P, Tosto G, Kushch N, Sha J, Bazemore K, et al. Multi-ancestry genome-wide meta-analysis of 56,241 individuals identifies LRRC4C, LHX5-AS1 and nominates ancestry-specific loci PTPRK , GRB14 , and KIAA0825 as novel risk loci for Alzheimer’s disease: the Alzheimer’s Disease Genetics Consortium. medRxiv : the preprint server for health sciences. 2023. doi:10.1101/2023.07.06.23292311

75. Salazar SV, Cox TO, Lee S, Brody AH, Chyung AS, Haas LT, et al. Alzheimer’s Disease Risk Factor Pyk2 Mediates Amyloid-β-Induced Synaptic Dysfunction and Loss. J Neurosci. 2019;39. doi:10.1523/JNEUROSCI.1873-18.2018

76. Giralt A, de Pins B, Cifuentes-Díaz C, López-Molina L, Farah AT, Tible M, et al. PTK2B/Pyk2 overexpression improves a mouse model of Alzheimer’s disease. Exp Neurol. 2018;307: 62–73.

77. Kilinc D, Vreulx A-C, Mendes T, Flaig A, Marques-Coelho D, Verschoore M, et al. Pyk2 overexpression in postsynaptic neurons blocks amyloid β1-42-induced synaptotoxicity in microfluidic co-cultures. Brain Commun. 2020;2: fcaa139.

78. Brody AH, Nies SH, Guan F, Smith LM, Mukherjee B, Salazar SA, et al. Alzheimer risk gene product Pyk2 suppresses tau phosphorylation and phenotypic effects of tauopathy. Mol Neurodegener. 2022;17: 1–33.

79. Foster EM, Dangla-Valls A, Lovestone S, Ribe EM, Buckley NJ. Clusterin in Alzheimer’s Disease: Mechanisms, Genetics, and Lessons From Other Pathologies. Front Neurosci. 2019;13: 164.

80. Humphreys DT, Carver JA, Easterbrook-Smith SB, Wilson MR. Clusterin has chaperone-like activity similar to that of small heat shock proteins. J Biol Chem. 1999;274: 6875–6881.

81. May PC, Lampert-Etchells M, Johnson SA, Poirier J, Masters JN, Finch CE. Dynamics of gene expression for a hippocampal glycoprotein elevated in Alzheimer’s disease and in response to experimental lesions in rat. Neuron. 1990;5: 831–839.

82. May PC, Robison P, Fuson K, Smalstig B, Stephenson D, Clemens JA. Sulfated glycoprotein-2 expression increases in rodent brain after transient global ischemia. Brain Res Mol Brain Res. 1992;15: 33–39.

83. Chaiwatanasirikul K-A, Sala A. The tumour-suppressive function of CLU is explained by its localisation and interaction with HSP60. Cell Death Dis. 2011;2: e219.

84. Merino-Zamorano C, Fernández-de Retana S, Montañola A, Batlle A, Saint-Pol J, Mysiorek C, et al. Modulation of Amyloid-β1-40 Transport by ApoA1 and ApoJ Across an in vitro Model of the Blood-Brain Barrier. J Alzheimers Dis. 2016;53: 677–691.

85. Yeh FL, Wang Y, Tom I, Gonzalez LC, Sheng M. TREM2 Binds to Apolipoproteins, Including APOE and CLU/APOJ, and Thereby Facilitates Uptake of Amyloid-Beta by Microglia. Neuron. 2016;91: 328–340.

86. Robbins JP, Perfect L, Ribe EM, Maresca M, Dangla-Valls A, Foster EM, et al. Clusterin Is Required for β-Amyloid Toxicity in Human iPSC-Derived Neurons. Front Neurosci. 2018;12: 504.

87. Killick R, Ribe EM, Al-Shawi R, Malik B, Hooper C, Fernandes C, et al. Clusterin regulates β-amyloid toxicity via Dickkopf-1-driven induction of the wnt-PCP-JNK pathway. Mol Psychiatry. 2014;19: 88–98.

88. Nilselid A-M, Davidsson P, Nägga K, Andreasen N, Fredman P, Blennow K. Clusterin in cerebrospinal fluid: analysis of carbohydrates and quantification of native and glycosylated forms. Neurochem Int. 2006;48: 718–728.

89. Nott A, Holtman IR, Coufal NG, Schlachetzki JCM, Yu M, Hu R, et al. Brain cell type-specific enhancer-promoter interactome maps and disease - risk association. Science. 2019;366. doi:10.1126/science.aay0793

90. Alasoo K, Rodrigues J, Mukhopadhyay S, Knights AJ, Mann AL, Kundu K, et al. Shared genetic effects on chromatin and gene expression indicate a role for enhancer priming in immune response. Nat Genet. 2018;50: 424–431.

91. Panousis NI, El Garwany O, Knights A, Rop JC, Kumasaka N, Imaz M, et al. Gene expression QTL mapping in stimulated iPSC-derived macrophages provides insights into common complex diseases. bioRxiv. 2023. p. 2023.05.29.542425. doi:10.1101/2023.05.29.542425

92. Tan M-S, Yang Y-X, Xu W, Wang H-F, Tan L, Zuo C-T, et al. Associations of Alzheimer’s disease risk variants with gene expression, amyloidosis, tauopathy, and neurodegeneration. Alzheimers Res Ther. 2021;13. doi:10.1186/s13195-020-00755-7

93. Bruntraeger M, Byrne M, Long K, Bassett AR. Editing the Genome of Human Induced Pluripotent Stem Cells Using CRISPR/Cas9 Ribonucleoprotein Complexes. Methods Mol Biol. 2019;1961: 153–183.

94. van Wilgenburg B, Browne C, Vowles J, Cowley SA. Efficient, long term production of monocyte-derived macrophages from human pluripotent stem cells under partly-defined and fully-defined conditions. PLoS One. 2013;8: e71098.

95. Douthwaite H, Arteagabeitia AB, Mukhopadhyay S. Differentiation of Human Induced Pluripotent Stem Cell into Macrophages. Bio Protoc. 2022;12: e4361.

96. Brownjohn PW, Smith J, Solanki R, Lohmann E, Houlden H, Hardy J, et al. Functional Studies of Missense TREM2 Mutations in Human Stem Cell-Derived Microglia. Stem Cell Reports. 2018;10: 1294–1307.

97. Washer SJ, Perez-Alcantara M, Chen Y, Steer J, James WS, Trynka G, et al. Single-cell transcriptomics defines an improved, validated monoculture protocol for differentiation of human iPSC to microglia. Sci Rep. 2022;12: 19454.

98. Haenseler W, Sansom SN, Buchrieser J, Newey SE, Moore CS, Nicholls FJ, et al. A Highly Efficient Human Pluripotent Stem Cell Microglia Model Displays a Neuronal-Co-culture-Specific Expression Profile and Inflammatory Response. Stem Cell Reports. 2017;8: 1727–1742.

99. Yu Y, Ye RD. Microglial Aβ receptors in Alzheimer’s disease. Cell Mol Neurobiol. 2015;35: 71–83.

100. Brandenburg L-O, Konrad M, Wruck CJ, Koch T, Lucius R, Pufe T. Functional and physical interactions between formyl-peptide-receptors and scavenger receptor MARCO and their involvement in amyloid beta 1-42-induced signal transduction in glial cells. J Neurochem. 2010;113: 749–760.

101. Ryan KJ, White CC, Patel K, Xu J, Olah M, Replogle JM, et al. A human microglia-like cellular model for assessing the effects of neurodegenerative disease gene variants. Sci Transl Med. 2017;9. doi:10.1126/scitranslmed.aai7635

102. Takaoka A, Tanaka N, Mitani Y, Miyazaki T, Fujii H, Sato M, et al. Protein tyrosine kinase Pyk2 mediates the Jak-dependent activation of MAPK and Stat1 in IFN-γ, but not IFN-α, signaling. EMBO J. 1999;18: 2480–2488.

103. El Khoury JB, Moore KJ, Means TK, Leung J, Terada K, Toft M, et al. CD36 Mediates the Innate Host Response to β-Amyloid. J Exp Med. 2003;197: 1657–1666.

104. Lee JK, Schuchman EH, Jin HK, Bae J-S. Soluble CCL5 Derived from Bone Marrow-Derived Mesenchymal Stem Cells and Activated by Amyloid β Ameliorates Alzheimer’s Disease in Mice by Recruiting Bone Marrow-Induced Microglia Immune Responses. Stem Cells. 2012;30: 1544–1555.

105. Zhang K, Tian L, Liu L, Feng Y, Dong Y-B, Li B, et al. CXCL1 Contributes to β-Amyloid-Induced Transendothelial Migration of Monocytes in Alzheimer’s Disease. PLoS One. 2013;8. doi:10.1371/journal.pone.0072744

106. Lee YK, Kwak DH, Oh KW, Nam S-Y, Lee BJ, Yun YW, et al. CCR5 deficiency induces astrocyte activation, Abeta deposit and impaired memory function. Neurobiol Learn Mem. 2009;92: 356–363.

107. Wojta KJ, Ayer AH, Ramos EM, Nguyen PD, Karydas AM, Yokoyama JS, et al. Lack of Association Between the CCR5-delta32 Polymorphism and Neurodegenerative Disorders. Alzheimer Dis Assoc Disord. 2020;34: 244–247.

108. Slaymaker IM, Gao L, Zetsche B, Scott DA, Yan WX, Zhang F. Rationally engineered Cas9 nucleases with improved specificity. Science. 2016;351: 84–88.

109. Li H, Durbin R. Fast and accurate long-read alignment with Burrows-Wheeler transform. Bioinformatics. 2010;26. doi:10.1093/bioinformatics/btp698

110. Ramírez F, Ryan DP, Grüning B, Bhardwaj V, Kilpert F, Richter AS, et al. deepTools2: a next generation web server for deep-sequencing data analysis. Nucleic Acids Res. 2016;44: W160–W165.

111. Liao Y, Smyth GK, Shi W. featureCounts: an efficient general purpose program for assigning sequence reads to genomic features. Bioinformatics. 2013;30: 923–930.

112. Ritchie ME, Phipson B, Wu D, Hu Y, Law CW, Shi W, et al. limma powers differential expression analyses for RNA-sequencing and microarray studies. Nucleic Acids Res. 2015;43: e47–e47.

113. Love MI, Huber W, Anders S. Moderated estimation of fold change and dispersion for RNA-seq data with DESeq2. Genome Biol. 2014;15: 1–21.

114. Robinson MD, McCarthy DJ, Smyth GK. edgeR: a Bioconductor package for differential expression analysis of digital gene expression data. Bioinformatics. 2009;26: 139–140.

115. Raudvere U, Kolberg L, Kuzmin I, Arak T, Adler P, Peterson H, et al. g:Profiler: a web server for functional enrichment analysis and conversions of gene lists (2019 update). Nucleic Acids Res. 2019;47: W191–W198.

116. Buenrostro J, Wu B, Chang H, Greenleaf W. ATAC-seq: A Method for Assaying Chromatin Accessibility Genome-Wide. Curr Protoc Mol Biol. 2015;109: 21.29.1.

117. Li H, Handsaker B, Wysoker A, Fennell T, Ruan J, Homer N, et al. The Sequence Alignment/Map format and SAMtools. Bioinformatics. 2009;25: 2078.

118. Zhang Y, Liu T, Meyer CA, Eeckhoute J, Johnson DS, Bernstein BE, et al. Model-based Analysis of ChIP-Seq (MACS). Genome Biol. 2008;9: 1–9.

119. Quinlan AR, Hall IM. BEDTools: a flexible suite of utilities for comparing genomic features. Bioinformatics. 2010;26: 841–842.

120. Yu G, Wang LG, He QY. ChIPseeker: an R/Bioconductor package for ChIP peak annotation, comparison and visualization. Bioinformatics. 2015;31. doi:10.1093/bioinformatics/btv145

121. Langmead B, Salzberg SL. Fast gapped-read alignment with Bowtie 2. Nat Methods. 9: 357.

122. Meers MP, Tenenbaum D, Henikoff S. Peak calling by Sparse Enrichment Analysis for CUT&RUN chromatin profiling. Epigenetics Chromatin. 2019;12: 1–11.

123. McQuin C, Goodman A, Chernyshev V, Kamentsky L, Cimini BA, Karhohs KW, et al. CellProfiler 3.0: Next-generation image processing for biology. PLoS Biol. 2018;16: e2005970.

124. Coetzee SG, Coetzee GA, Hazelett DJ. motifbreakR: an R/Bioconductor package for predicting variant effects at transcription factor binding sites. Bioinformatics. 2015;31: 3847.

