## Supplementary material for "An Alzheimer’s disease-associated common regulatory variant in a PTK2B intron alters microglial function": Supp Fig

**Consisting of**
Supplementary figures 1-5
Supplementary table legends
Supplementary references

### Supplementary Figures

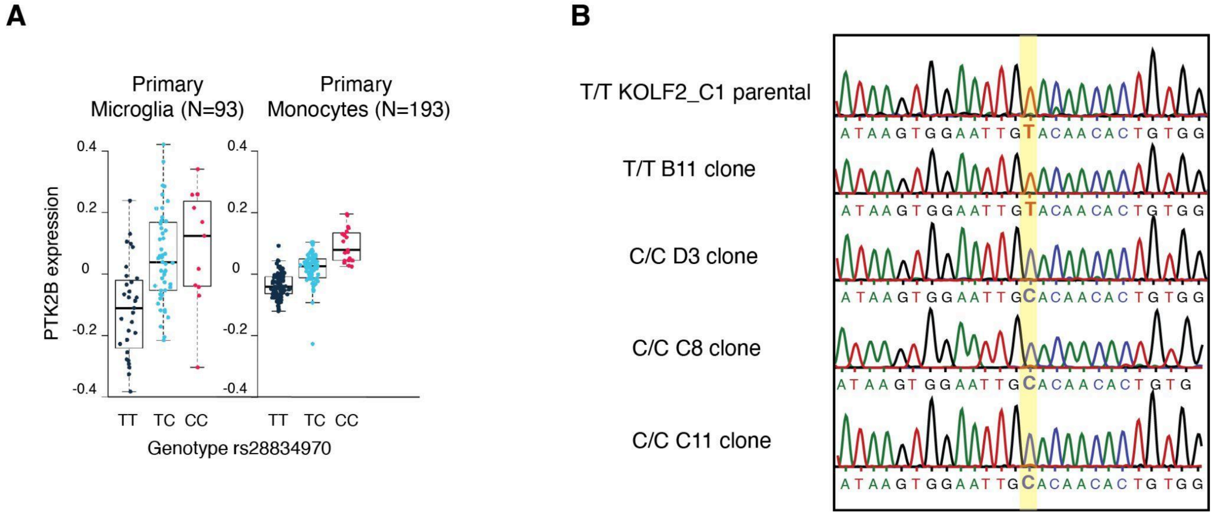


**Supp Figure 1 - Characterisation of the rs28834970 variant in PTK2B**

A) Boxplots showing the expression of the *PTK2B* gene stratified by the rs28834970 genotype in primary microglia and primary monocytes from[25]. The y-axis shows normalised expression levels (log TPM value) and each dot on the box shows the expression level of a single sample.

B) Sanger sequencing tracks of an amplicon around the rs28834970 variant in wild-type T/T KOLF2_C1 hiPSC clones (the KOLF2_C1 parental line and B11 T/T clone which has been through the editing and clonal selection process) and engineered homozygous C/C KOLF2_C1 clones (D3, C8 and C11). The rs28834970 variant is highlighted in yellow.

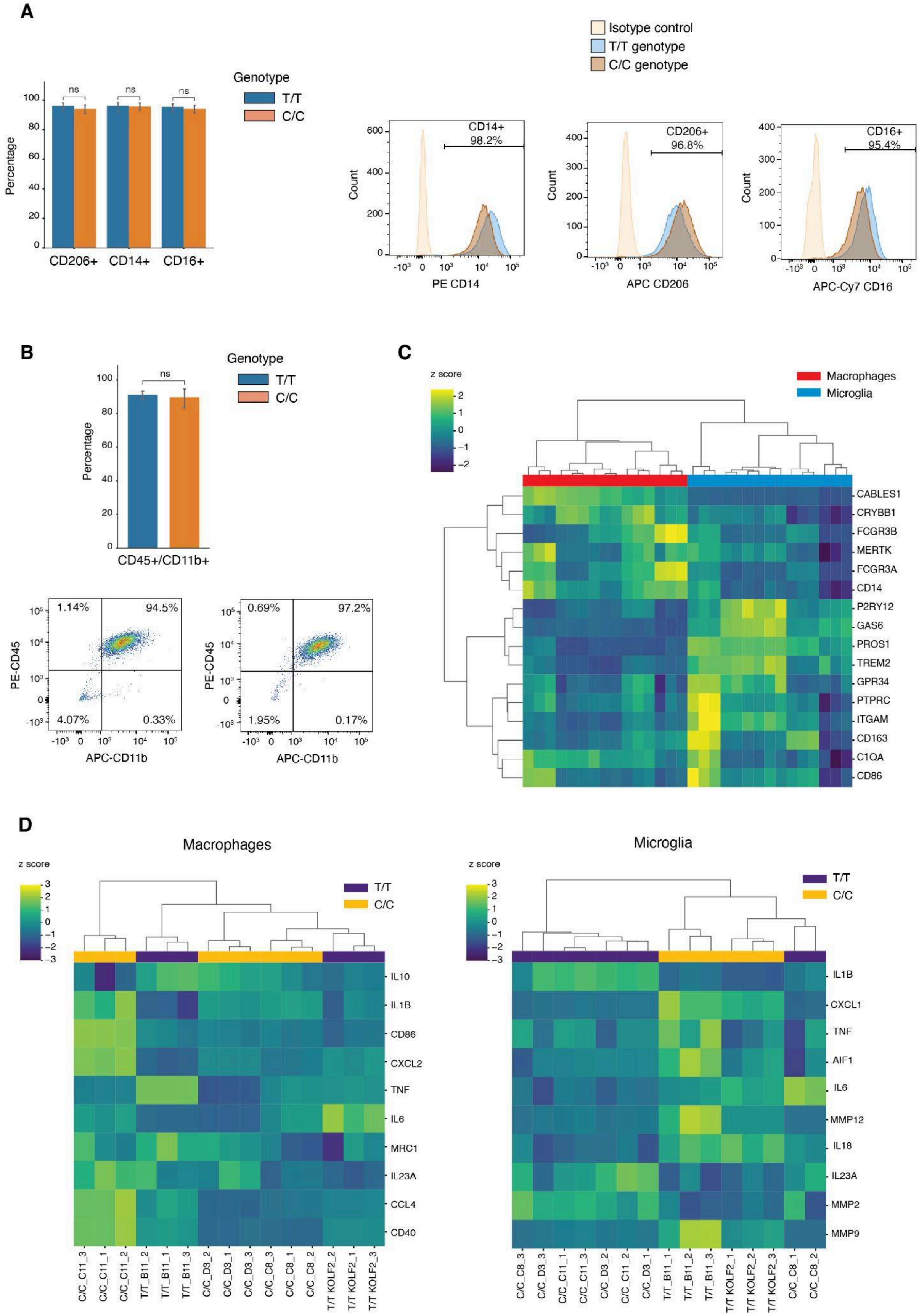


**Supp Figure 2 - Characterisation of differentiation of hiPSC-derived macrophages and microglia**

A) Bar graph showing the percentage of macrophages harbouring either the T/T or C/C allele positive for each marker (left). Results are shown as the mean ± SEM of independent biological replicates (n=3). ns= non-significant, unpaired t-test. Representative histograms from FACS analysis of single-stained macrophage markers CD14, CD16 and CD206 in hiPSC-derived macrophages with either T/T or C/C allele at rs28834970 (right).
B) Bar graph showing the percentage of CD11b+CD45+ microglia with the T/T or C/C allele (top). Results are shown as the mean ± SEM of independent biological replicates (n=3). ns= non-significant, unpaired t-test. Representative scatterplots from FACS analysis of microglia markers CD11b and CD45 in hiPSC-derived microglia harbouring the T/T or C/C allele at rs28834970 (bottom).
C) Heatmap showing the relative expression of microglia and macrophage markers[75,101,102] in hiPSC-derived macrophages (red) and microglia (blue) harbouring the T/T or C/C allele at rs28834970.
D) Heatmap showing the relative expression of activation markers[103,104] of macrophages (left) and microglia (right) in the corresponding hiPSC-derived cell type harbouring either the T/T (orange) or the C/C (purple) allele at rs28834970.

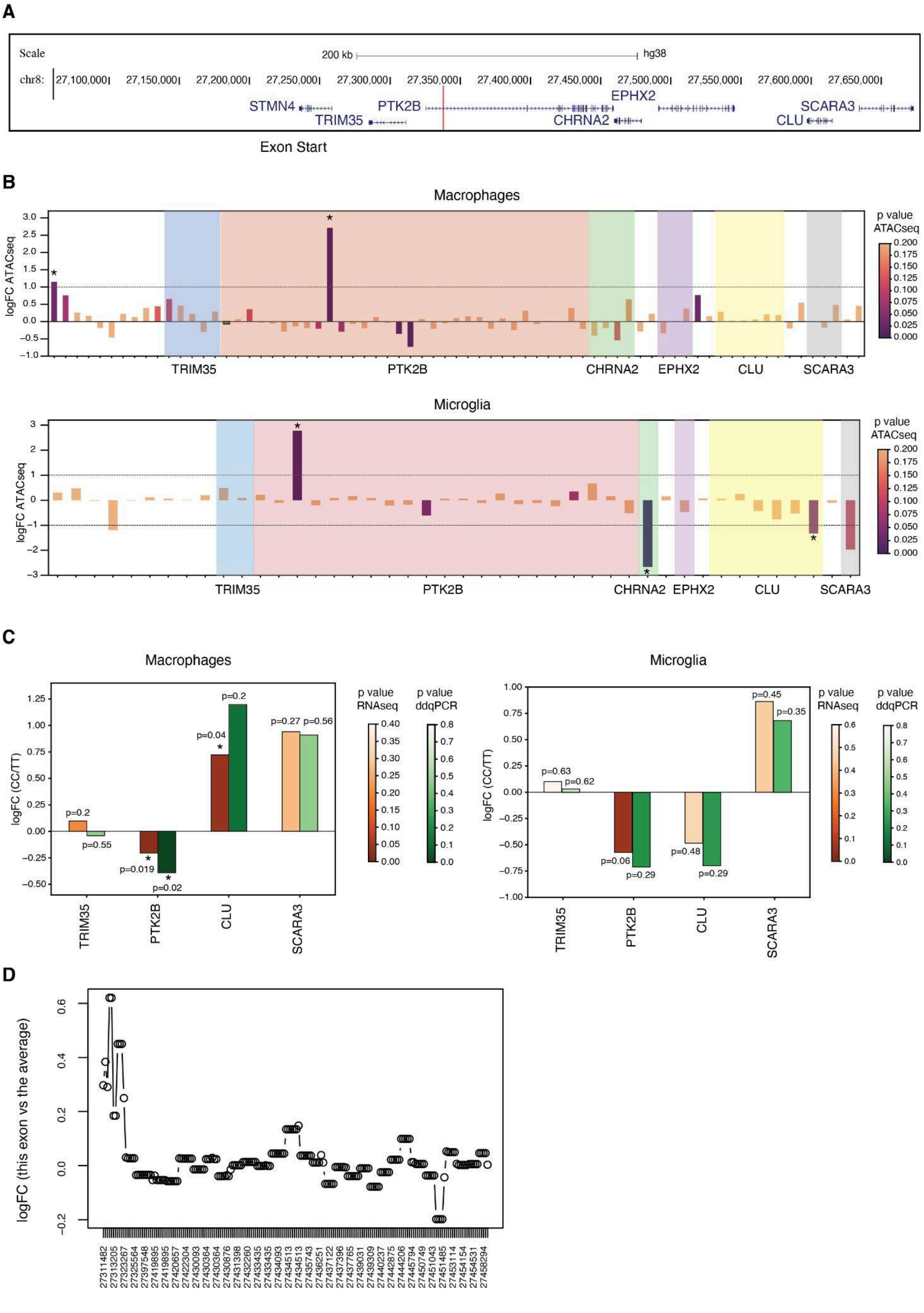


**Supp Figure 3 - Effects of the rs28834970 variant on the surrounding genomic region**

A) Track from UCSC Genome Browser (Human GRCh38/hg38) showing RefSeq genes in a 600 kb window around the rs28834970 SNP (highlighted in red).

B) Bar graph showing logFC of chromatin accessibility between macrophages (top) and microglia (bottom) harbouring the C/C versus the T/T allele for all ATAC-seq peaks identified in the 600kb window around rs28834970. The colour of the bars indicates the p value and peaks (bars) located in the same gene are highlighted in the same colour. The asterisks denote peaks reaching significance (logFC>+-1 and $p<0.05$).

C) Bar graph showing the log fold change (FC) of expression of the genes located in the window around rs28834970 in macrophages (left) and microglia (right) with the C/C versus the T/T allele, measured by RNAseq (red) and ddqPCR (green). ddqPCR n=3 * $p<0.05$, **$p<0.01$ unpaired t-test.

D) Plot showing the results of the analysis of differential exon usage between microglia harbouring the C/C and the T/T allele at rs28834970. The relative logFC of expression of each exon of *PTK2B* is the difference between the exon's logFC and the overall logFC for the gene. None of the *PTK2B* exons (shown as dots) were significantly changed.

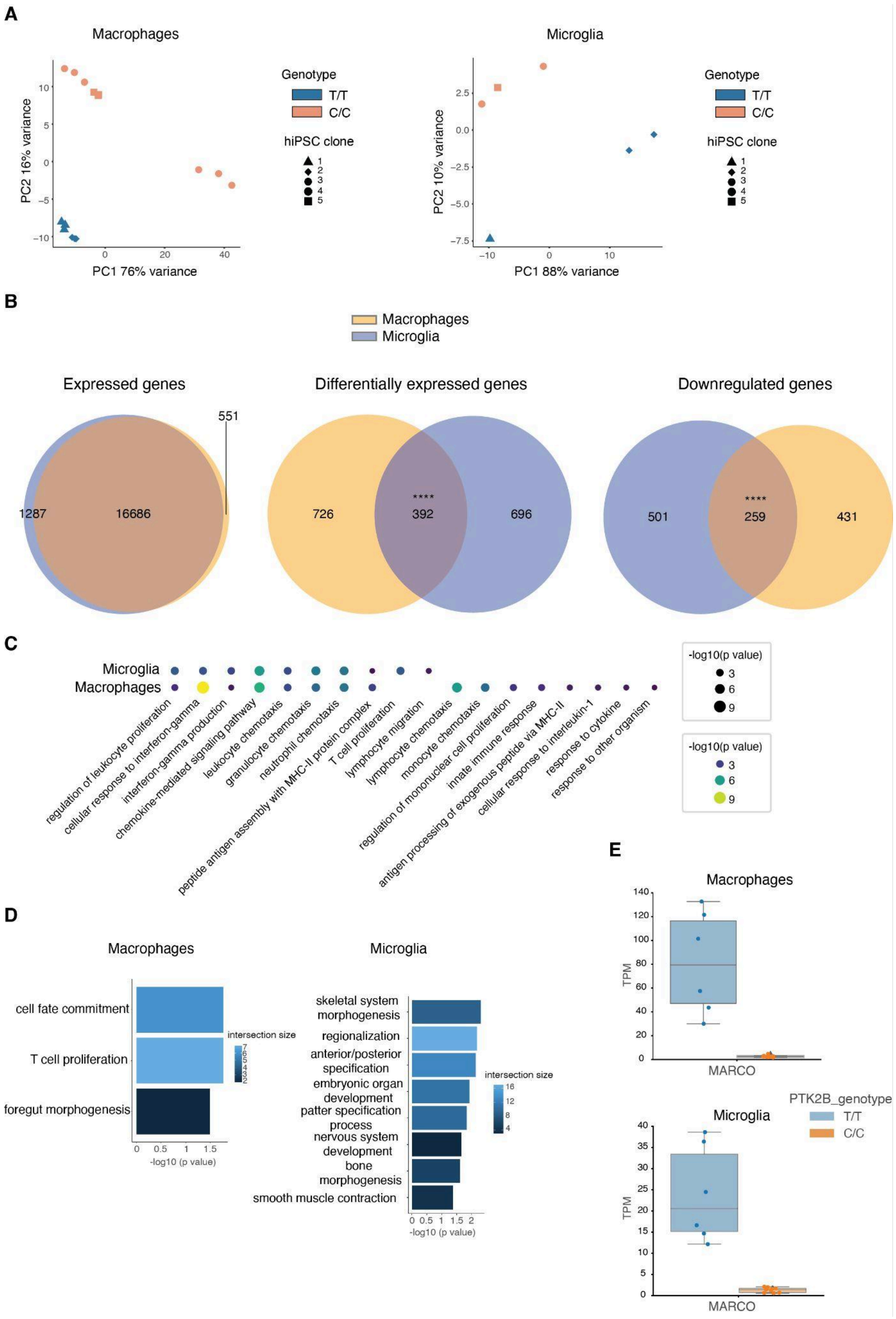
A
Macrophages
Microglia
PC2 16% variance
PC1 76% variance
PC2 10% variance
PC1 88% variance
Genotype
T/T
C/C
hiPSC clone
1
2
3
4
5
B
Macrophages
Microglia
Expressed genes
551
1287
16686
Differentially expressed genes
726
****
392
696
Downregulated genes
501
****
259
431
C
Microglia
Macrophages
regulation of leukocyte proliferation
cellular response to interferon-gamma
interferon-gamma production
chemokine-mediated signaling pathway
leukocyte chemotaxis
granulocyte chemotaxis
neutrophil chemotaxis
peptide antigen assembly with MHC-II protein complex
T cell proliferation
lymphocyte migration
lymphocyte chemotaxis
monocyte chemotaxis
regulation of mononuclear cell proliferation
innate immune response
antigen processing of exogenous peptide via MHC-II
cellular response to interleukin-1
response to cytokine
response to other organism
-log10(p value)
D
Macrophages
cell fate commitment
T cell proliferation
foregut morphogenesis
intersection size
-log10 (p value)
Microglia
skeletal system morphogenesis
regionalization
anterior/posterior specification
embryonic organ development
patter specification process
nervous system development
bone morphogenesis
smooth muscle contraction
intersection size
-log10 (p value)
E
Macrophages
TPM
MARCO
Microglia
TPM
MARCO
PTK2B_genotype
T/T
C/C

**Supp Figure 4 - Characterisation of the effects of rs28834970 on chromatin accessibility and gene expression in iPSC-derived macrophages and microglia**

A) Principal component analysis (PCA) of accessible regions across the genome of macrophages (left) and microglia (right) differentiated from two hiPSC clones with the T/T allele and three clones with the C/C allele, measured by ATAC-seq.

B) Venn diagrams showing overlaps between microglia and macrophages of all expressed genes (left), all differentially expressed genes (middle) and downregulated genes (logFC<-0.5 and p<0.05) (right), measured by RNA-seq. Statistical significance of number of overlaps was calculated using Chi-square test with Yates' correction. ****p<0.0001

C) Top pathways identified in GO analysis of downregulated genes in microglia and macrophages. Colour and size of the dots represents the -log10 of the p-value.

D) Results of GO analysis of upregulated genes (logFC>0.5 and p<0.05) in macrophages (left) and microglia (right) with the C/C versus the T/T allele at rs28834970.

E) Expression of *MARCO* in macrophages (top) and microglia (bottom) harbouring the T/T or C/C allele at rs28834970, measured by RNA-seq. Cells were differentiated from two hiPSC clones with the T/T allele and three clones with the C/C allele. TPM= transcripts per million, differential expression analysis performed using DESeq2.

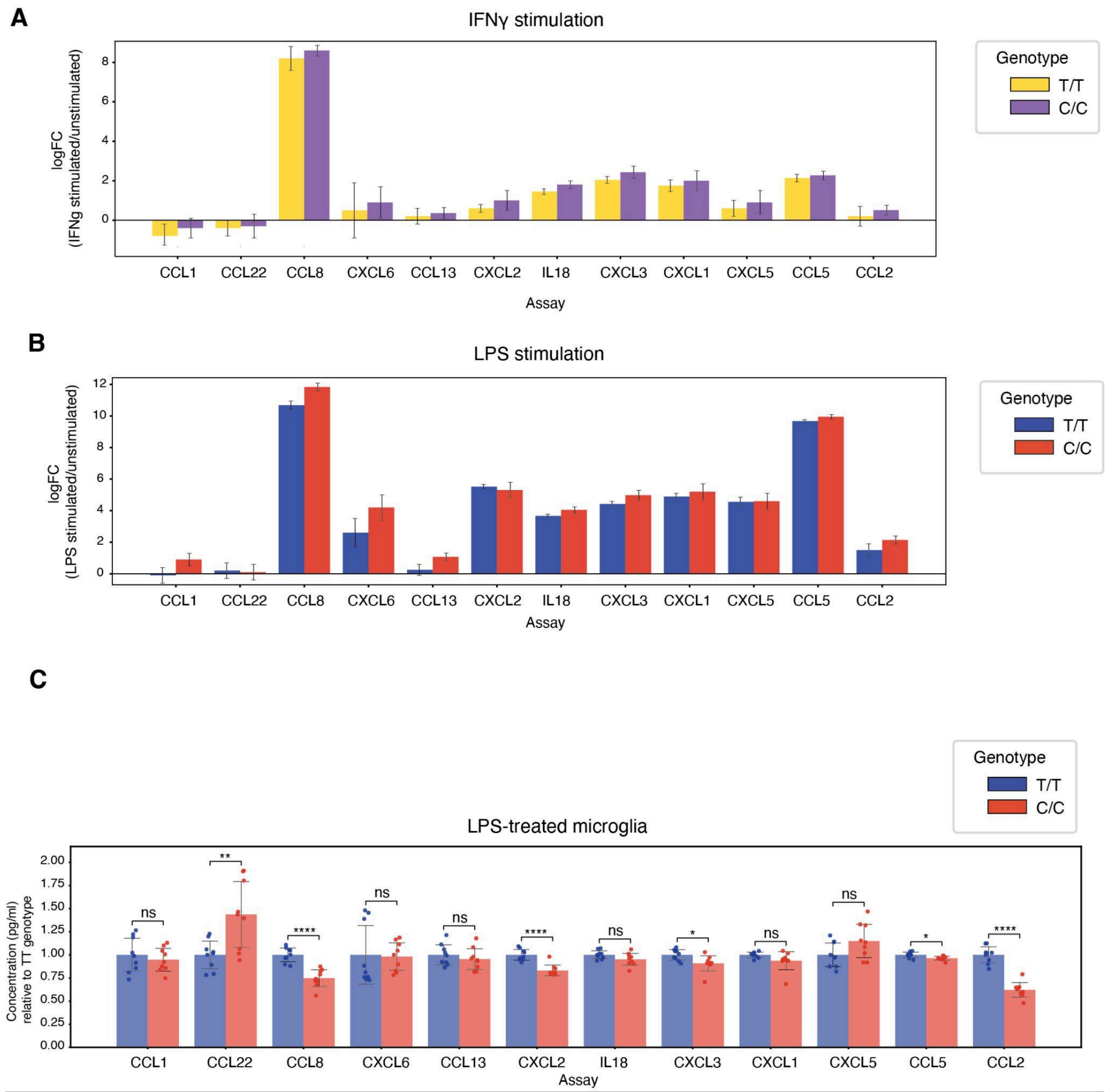


**Supp Figure 5 - The rs28834970 variant causes changes in chemokine production iPSC-derived microglia**

A) Bar graph showing logFC of chemokine concentration between unstimulated microglia and microglia stimulated with IFNγ. Yellow bars represent microglia with the T/T allele and purple bars show logFC of microglia with the C/C allele. Results are shown as the mean ± propagation of error of independent biological replicates (n=3).

B) Bar graph showing logFC of chemokine concentration between unstimulated microglia and microglia stimulated with LPS. Blue bars represent microglia with the T/T allele and red bars show logFC of microglia with the C/C allele. Results are shown as the mean ± propagation of error of independent biological replicates (n=3).

C) Concentration of downregulated chemokines in microglia with the C/C allele at rs28834970 relative to the concentration in cells with the T/T allele, stimulated with LPS. Results are shown as the mean ± SEM of independent biological replicates (n=3). 3 technical replicates of each were measured. ns= non-significant, * p<0.05, **p<0.01, ***p<0.001, ****p<0.0001, unpaired t-test.

### Supplementary Table Legends

**Supp Table 1 - ATACseq analysis in macrophages and microglia**
Summary of results for all ATACseq peaks in macrophages or microglia. Table shows peak position, mean counts (baseMean), fold changes in C/C relative to T/T (log2FoldChange) with its respective standard error (lfcSE), the Wald statistics (log2FoldChange divided by lfcSE) (stat) and p-values before (pvalue) and after (padj) multiple testing correction.

**Supp Table 2 - CEBPβ CUT&RUN analysis in microglia**
Summary of results for all CEBPβ CUT&RUN peaks in microglia. Table shows peak position, mean counts (baseMean), fold changes in C/C relative to T/T (log2FoldChange) with its respective standard error (lfcSE), the Wald statistics (log2FoldChange divided by lfcSE) (stat) and p-values before (pvalue) and after (padj) multiple testing correction.

**Supp Table 3 - RNAseq analysis in macrophages and microglia**
Summary of results of RNAseq analysis in macrophages or microglia. Table shows Ensembl gene ID (ENSEMBL), gene symbol (SYMBOL), full gene name (GENENAME), basal expression (baseMean), fold changes in C/C relative to T/T (log2FoldChange) with its respective standard error (lfcSE), the Wald statistics (log2FoldChange divided by lfcSE) (stat) and p-values before (pvalue) and after (padj) multiple testing correction.

**Supp Table 4 - ddqPCR assay in macrophages and microglia**
Summary of results of ddqPCR assay in macrophages or microglia. Table shows raw values of the relative concentration of each target gene (copies/ul relative to copies/ul of housekeeping gene) in cells with T/T or C/C genotype.

**Supp Table 5 - Luminex assay in microglia**
Summary of results of Luminex assay in microglia. Table shows concentration (pg/ml) of each chemokine measured in microglia with T/T or C/C genotype.

**Supp Table 6 - ENA accession numbers for the datasets**
Summary of the ENA accession numbers (SampleID) and linkage to the respective sequencing samples (Sample Name).

### Supplementary references

1. Young, A. M. H. *et al.* A map of transcriptional heterogeneity and regulatory variation in human microglia. *Nat. Genet.* **53**, (2021).
2. Brownjohn, P. W. *et al.* Functional Studies of Missense TREM2 Mutations in Human Stem Cell-Derived Microglia. *Stem Cell Reports* **10**, 1294–1307 (2018).
3. Haenseler, W. *et al.* A Highly Efficient Human Pluripotent Stem Cell Microglia Model Displays a

Neuronal-Co-culture-Specific Expression Profile and Inflammatory Response. *Stem Cell Reports* **8**, 1727–1742 (2017).

4. Vaughan-Jackson, A. *et al.* Differentiation of human induced pluripotent stem cells to authentic macrophages using a defined, serum-free, open-source medium. *Stem Cell Reports* **16**, 3093 (2021).
5. Jurga, A. M., Paleczna, M. & Kuter, K. Z. Overview of General and Discriminating Markers of Differential Microglia Phenotypes. *Front. Cell. Neurosci.* **14**, 544457 (2020).
6. Murray, P. J. *et al.* Macrophage activation and polarization: nomenclature and experimental guidelines. *Immunity* **41**, (2014).
